# Macrophage vacuolar ATPase (v-ATPase) function controls *Aspergillus fumigatus* germination and hyphal growth independent of spore killing

**DOI:** 10.1101/2025.07.14.664761

**Authors:** Aqib R. Magray, Emily E. Rosowski

## Abstract

Tissue-resident macrophages efficiently internalize *Aspergillus fumigatus* spores, forming a critical first line of defense against infection. However, the mechanisms that these cells use to control spores in vivo remain incompletely defined. Here, we used the live imaging capabilities of the larval zebrafish host model to assess the role of the v-ATPase complex in macrophage-mediated defense against *A. fumigatus* in a whole vertebrate animal. For the first time we are able to visualize co-localization of *A. fumigatus* spores with the key v-ATPase subunit Atp6v1h in macrophages inside of an infected animal. As macrophages only have a low ability to kill spores, this co-localization occurs as early as 1-day post-injection and persists for multiple days. Surprisingly, macrophage spore killing is not further reduced by targeting of *atp6v1h* with CRISPR/Cas9. Instead, v-ATPase deficiency profoundly impacts macrophage-mediated control of spore swelling, decoupling the macrophage functions of spore killing and inhibition of germination. We also identify a role for the v-ATPase complex in macrophage control of extracellular hyphal growth. These effects on macrophage function drive significantly decreased host survival in larvae lacking a functional v-ATPase. We also report broad effects of v-ATPase deficiency on macrophage numbers, apoptosis in the hematopoietic tissue, and potential neutrophil functions, reflecting the importance of this complex in host antifungal immunity.

**Graphical Abstract:** 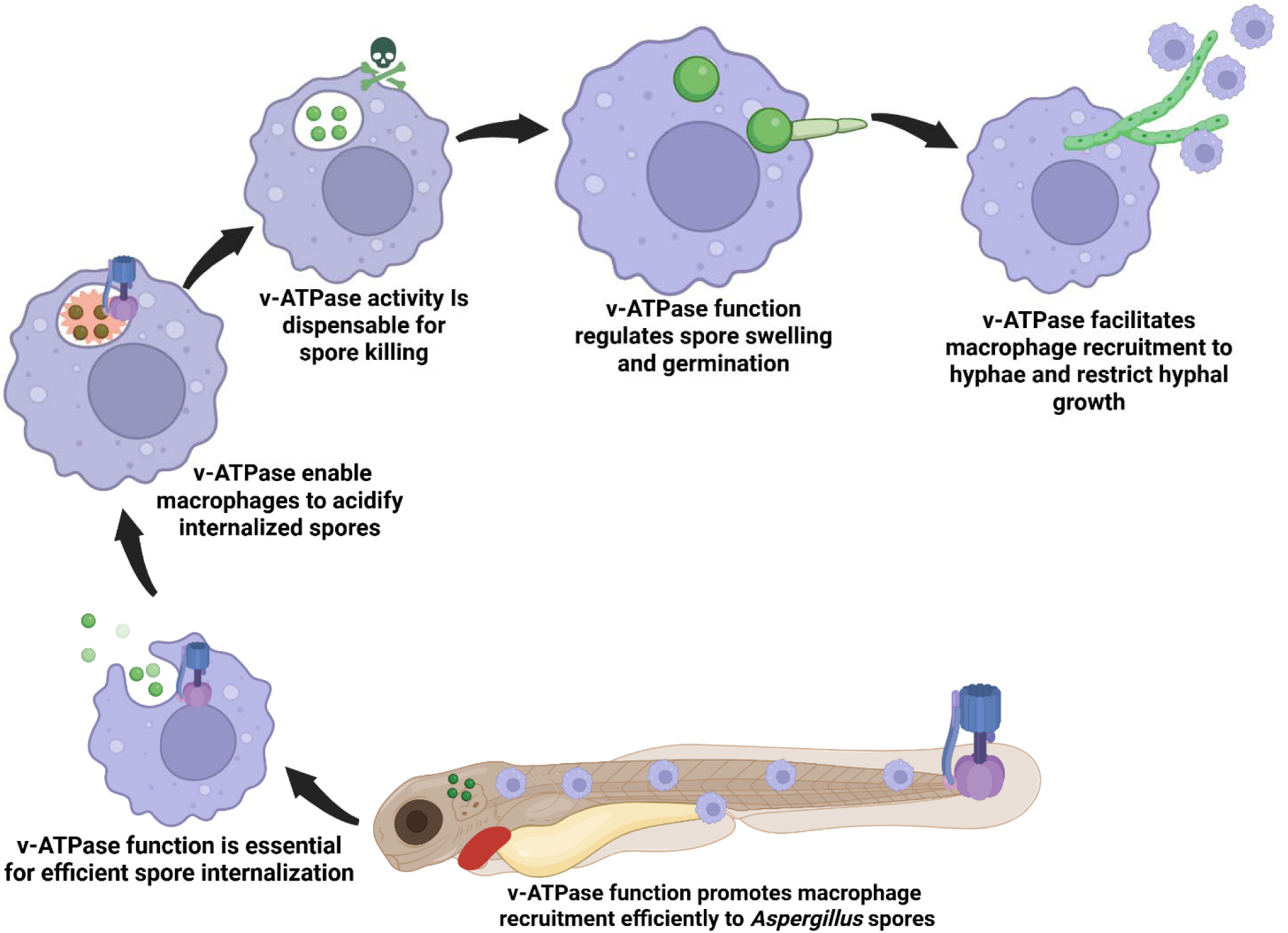

**Highlights:**

- **v-ATPase function in macrophages restricts *Aspergillus* germination and hyphal growth**
- **Spore killing by macrophages is independent of v-ATPase function**
- **Loss of macrophage v-ATPase mimics the effects of macrophage deficiency in vivo**

## Introduction

*Aspergillus fumigatus*, a ubiquitous airborne fungus, causes Invasive Pulmonary Aspergillosis (IPA) primarily in immunocompromised individuals, hematopoietic stem cell transplant recipients, structural lung disease patients and corticosteroid users^1–5^. In immunocompetent hosts, inhaled *A. fumigatus* spores are efficiently cleared by mucociliary mechanisms, taken up by epithelial cells, or phagocytosed by alveolar macrophages, the latter of which inhibit spore germination and prevent invasive hyphal growth ^6–8^. Macrophages constitute 80% of immune cells in healthy lungs and are therefore the principal resident phagocytic cell against inhaled fungal spores (conidia) within alveoli ^9^. However, the ability of macrophages to completely eradicate *A. fumigatus* spores in vivo is limited. For example, studies in larval zebrafish show that nearly 50% of engulfed spores can persist within macrophages for up to seven days ^8^. In mice, it can take up to 3 days for alveolar macrophages to eradicate spores ^10^ and neutrophil recruitment is generally required for host survival ^9^. *A. fumigatus* employs fungal factors that subvert host defenses by interfering with immune recognition and hijacking phagolysosomal pathways, thereby promoting spore survival and hyphal formation ^11–15^.

One of the key antimicrobial strategies of macrophages involves phagolysosomal acidification, a process critically dependent on the vacuolar-type ATPase (v-ATPase) complex^16,17^.

This evolutionarily conserved, ATP-driven proton pump comprises a cytosolic V1 domain and a membrane-bound Vo domain, which work in tandem to acidify intracellular compartments such as late endosomes and lysosomes. The resulting low pH is essential for proper enzyme trafficking and the activation of lysosomal hydrolases, thereby leading to efficient pathogen degradation ^18–21^. In phagocytes, including macrophages and neutrophils, inhibition of v-ATPase-mediated acidification impairs microbial killing, including of the fungi *Candida albicans* and *Cryptococcus neoformans* ^22–25^. Previous studies on *A. fumigatus* infection of alveolar macrophages in cell culture also suggests that *A. fumigatus* spore killing is inhibited by treatment with the v-ATPase inhibitor bafilomycin A1 ^26,27^. However, the role of v-ATPase-driven phagolysosomal acidification in overall macrophage-mediated fungal control in a whole animal in vivo model and its impact on host survival has not been directly investigated.

Here, we address this knowledge gap by using a larval zebrafish infection model to dissect the functional importance of macrophage acidification machinery in antifungal innate immunity. In larval zebrafish, macrophages and neutrophils constitute the principal immune effector cells, as the adaptive immune system remains immature until 4–6 weeks post-fertilization, allowing for the exclusive study of innate immune responses ^28^. Additionally, transgenic neutrophil-defective models allow for the specific investigation of macrophage-mediated control of infection ^29^. The larval zebrafish–*Aspergillus* infection model is well-established and has yielded multiple insights into innate immunity to this pathogen ^8,30–32^.

We report that fluorescently-tagged Atp6v1h, the essential H subunit required for v-ATPase assembly and proton-pumping activity, persistently co-localizes with 35-50% of macrophage-phagocytosed spores in larval zebrafish across 5 days of infection. Neutrophil-defective *atp6v1h* crispant larvae have increased susceptibility to *A. fumigatus*, demonstrating the importance of the v-ATPase complex for macrophage-mediated control of infection. This susceptibility is partially due to an overall decrease in macrophage number across whole larvae, suggesting that the v-ATPase complex is required for macrophage health and development. Focusing on macrophages that migrate to the infection site and take up spores, we confirm that v-ATPase deficiency results in decreased acidification of phagocytosed spores. Surprisingly, however, we find no significant difference in macrophage-mediated spore killing in *atp6v1h* crispant larvae. Daily imaging of fungal development inside of larvae reveals that v-ATPase activity promotes macrophage-mediated control of spore swelling without affecting spore killing. These results therefore demonstrate that control of spore germination can be uncoupled with spore killing. Imaging of larvae after fungal spore germination also reveals that v-ATPase activity promotes macrophage-mediated control of hyphal growth. Altogether, our results highlight the importance of the v-ATPase complex for macrophage function against pathogens in vivo while establishing that this pathway is not required for direct *A. fumigatus* spore killing.

## Material and Methods

### Zebrafish lines and husbandry

Zebrafish lines, embryos, and larvae were maintained and handled according to protocols approved by the Clemson University Institutional Animal Care and Use Committee (IACUC) (AUP2021-0109, AUP2022-0093, and AUP2022-0111), following the Guide for the Care and Use of Laboratory Animals and complying with the regulations and guidelines set forth in the Animal Welfare Act and Regulations. Adult zebrafish were housed at 28°C on a 14-hour light/10-hour dark cycle and were fed twice a day. All transgenic and mutant fish lines used in this study were maintained in the AB genetic background and are listed in Table 1. Zebrafish adults and larvae were maintained under standard experimental conditions ^33,34^. All embryos and larvae were anesthetized in 0.3 mg/mL buffered tricaine (MS-222) prior to any experimental manipulations. The *irf8* mutant line was maintained as heterozygotes by out-crossing and adult *irf8*^+/-^ fish were in-crossed to generate *irf8*^+/+^, *irf8*^+/-^, and *irf8*^-/-^ progeny that were genotyped at the conclusion of the experiment as previously described ^35^.

**Table 1.** Zebrafish lines used in this study.

| Zebrafish line | Purpose | Fluorescence | References |
| --- | --- | --- | --- |
| Tg(*mpx:mCherry-2A-rac2D57N*) | Neutrophil-defective | mCherry in neutrophil cytoplasm | <sup>38</sup> |
| Tg(*lyz:mCherry-H2B*) | Label neutrophil nuclei | mCherry in neutrophil nuclei | <sup>33</sup> |
| Tg(*mpeg1:GFP-H2B*) | Label macrophage nuclei | GFP in macrophage nuclei | <sup>86</sup> |
| Tg(*mfap4:BFP*) | Label macrophage cytoplasm | BFP in macrophage cytoplasm | <sup>34</sup> |
| *irf8*<sup>-/-</sup> | Macrophage-defective | No fluorescence | <sup>59</sup> |
| Tg(*mfap4:TagRFPT.zf1-atp6v1h_cyaa-mCherry*) | Express fluorescently-tagged Atp6v1h under macrophage-specific promoter | RFP-tagged Atp6v1h in macrophages and mCherry in eye | This study |

### Generation of fluorescently-tagged atp6v1h transgenic zebrafish

First, a *TagRFPT* sequence codon-optimized for expression in zebrafish was amplified from pCS2-TagRFPT.zf1 ^36^ (Addgene 61390, a gift from Harold Burgess) with Q5 polymerase (NEB) without the stop codon and adding a sequence encoding a 7 amino acid glycine-serine linker including a BamHI site at the C terminal end (Table 2). This was cloned into the SalI site of Tol2-*mfap4* ^37^ (Addgene 232189) by HiFi cloning (NEB). Coding sequence of *atp6v1h* was amplified from cDNA made with iScript RT supermix (Bio-Rad) from total RNA isolated from 2-3 dpf zebrafish larvae with TRIzol (Invitrogen) and cloned into the BamHI site directly after *TagRFPT* (Table 2). RefSeq predicts two splicing isoforms of *atp6v1h* with either 14 total exons (including non-coding) or 13 total exons (missing exon 7). Amplified sequence contained all 14 exons and was Sanger sequenced to confirm that it matches transcript NM_001291708. A *cyraa* promoter driving *mCherry* expression in the lens of the eye was also cloned into the MfeI site on the reverse strand, as a marker for genomic integration. Coding sequence for *mCherry* was amplified from Tol2-*mpx*:*mCherry-2A-rac2* ^38^ (a gift from Anna Huttenlocher) and sequence of the *cryaa* promoter was amplified from hsp70l-loxP-mCherry-STOP-loxP-H2B-GFP_cryaa-cerulean ^39^ (Addgene 24334, a gift from Didier Stainier) and these fragments were cloned sequentially into the Tol2 backbone by HiFi cloning (NEB)(Table 2).

**Table 2:**
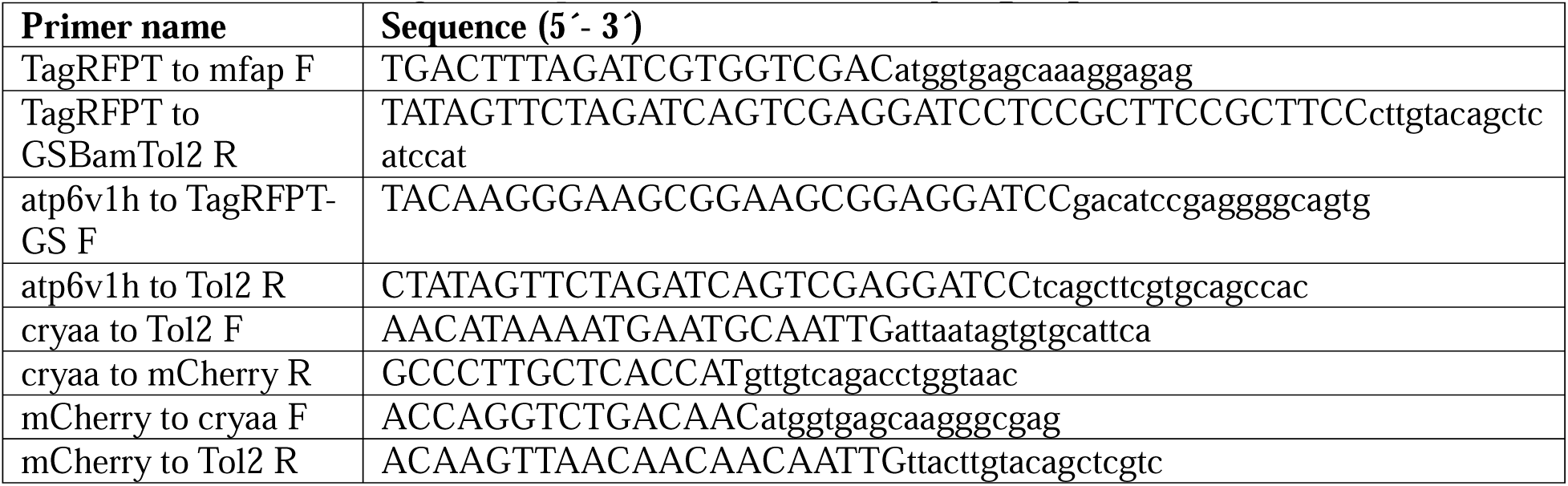
Primers used to generate plasmid to fluorescently-tag Atp6v1h.

For integration of Tol2-*mfap4:TagRFPT.zf1-atp6v1h cyraa:mCherry* into the zebrafish genome, *Tol2 transposase* was in vitro transcribed from NotI-digested pCS2-*transposase* (a gift from Anna Huttenlocher) using an mMESSAGE mMACHINE SP6 kit (Invitrogen) and mRNA was purified with a MEGAclear kit (Invitrogen) according to manufacturers’ instructions. Then, 1-2 nl of an injection mix containing 20 ng/ml plasmid and 10 ng/ml *transposase* mRNA was injected into the yolk of single-cell embryos of the AB strain. Injected F0 embryos were grown to adulthood, and a founder with integration of the DNA into the germline was determined by outcrossing single F0 adults and screening for mCherry and TagRFPT expression.

### CRISPR gRNA design, injection, and TIDE analysis

Guide RNAs (gRNAs) were designed using the CHOPCHOP web tool (https://chopchop.cbu.uib.no/) ^40,41^ (Suppl. Fig 1A, Table 3). gRNAs were designed to target exons 3 and 5 of the *atp6v1h* gene. These exons encode highly conserved motifs that are critical for catalytic activity and maintaining the integrity of the protein complex. Importantly, the targeted sequences correspond to essential domains annotated in the human homolog on UniProt (UniProt Consortium, 2023) and are conserved between humans and zebrafish. gRNAs targeting *luciferase* coding sequence were used as a control ^37^. To in vitro synthesize gRNAs, DNA oligos were used to create a double-stranded DNA template containing a T7 promoter sequence (5’-TAATACGACTCACTATAG -3’), the gRNA target sequence without the PAM, and the Cas9 scaffold sequence (5’-GTTTTAGAGCTAGAAATAGCAAGTTAAAATAAGGCTAGTCCGTTATCAACTTGAAAAAGTGGCACCGAGTCGGTGCTTTT -3’) (Suppl. Fig 1B). The template was then column-purified, and in vitro transcription was performed using the HiScribe T7 High Yield RNA Synthesis Kit (NEB). Template DNA was removed using DNase I (NEB), and the gRNAs were purified using the Monarch RNA Cleanup Kit (NEB), aliquoted, and stored at -80°C. A microinjection setup (BTX, Microject 1000A) equipped with a pressure injector, micromanipulator (Narishige), micropipette holder, footswitch, and compressed nitrogen gas was used to inject approximately ∼2 nl of a mixture containing 100 ng/µl of each gRNA, 250 ng/ml of Cas9 protein with an NLS resuspended in deionized (DI) water (PNAbio), and 0.25% phenol red into the yolk of 1 cell stage naturally spawned embryos.

**Table 3:**
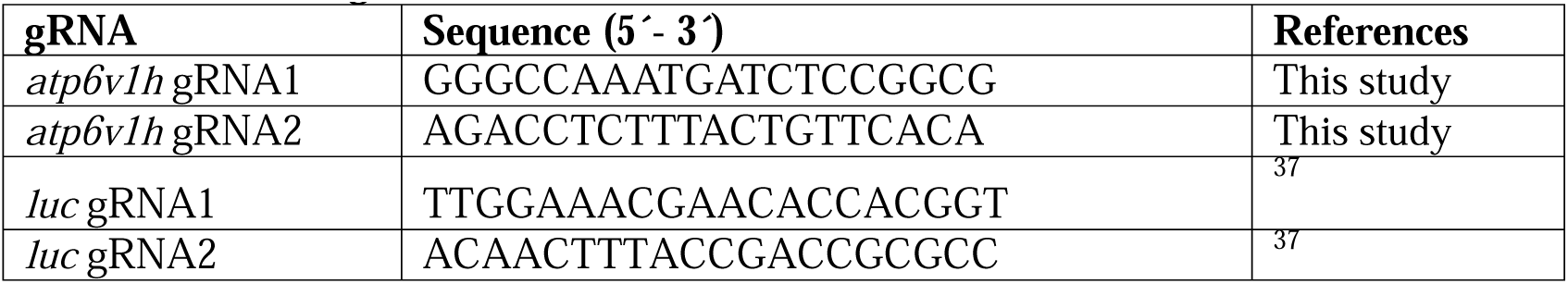
CRISPR gRNAs.

To evaluate the efficiency of gRNA-mediated gene disruption, PCR and gel electrophoresis as well as TIDE analysis was done. First, primers were designed to directly flank the gRNA target sites (F1+R1, F2+R2) (Table 4, Suppl. Fig 1A). PCR was performed from extracted genomic DNA from 1-2 dpf larvae using GoTaq polymerase and run on a 2.5% agarose gel. Amplification of each target sequence shows smearing demonstrating gene disruption due to indels generated because of DNA repair (Suppl. Fig 1C). Additionally, amplification of the region containing both target sites demonstrates that the entire DNA sequence between target sites can also be lost (Suppl. Fig 1C). To quantify gRNA efficiency with the TIDE web tool (https://tide.nki.nl/) ^42^, primers were designed to amplify 400-500 bp around each target site for gRNA1 and gRNA2 with primers sets, *atp6v1h*_gF1_TIDE; *atp6v1h*_gR1_TIDE, and *atp6v1h*_gF2_TIDE; *atp6v1h*_gF2_TIDE respectively (Table 4, Suppl. Fig 1A). PCR products were Sanger sequenced with both forward and reverse primers (Eton Bioscience) and TIDE analysis demonstrates efficiencies of ∼55% and ∼68% for gRNAs 1+2, respectively (Suppl. Fig 1D).

**Table 4:**
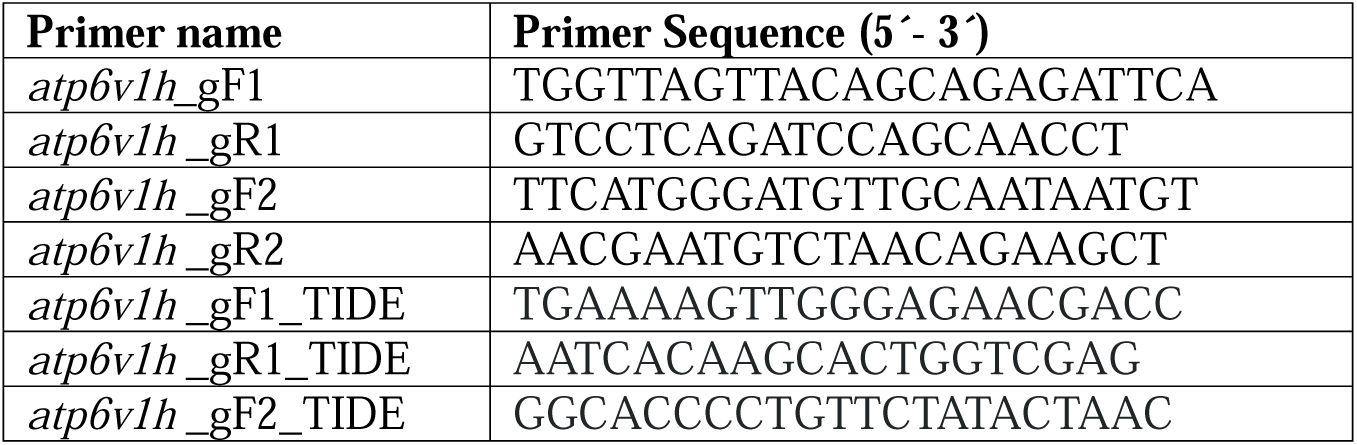
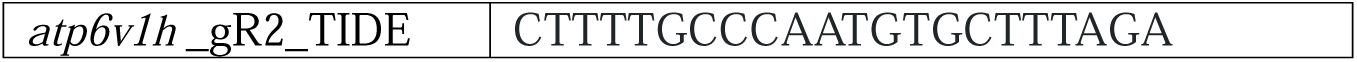
Primers used to test successful alterations of DNA.

### *Aspergillus fumigatus* strains and preparation of spores

All *A. fumigatus* strains used in this study are detailed in Table 5, including *Af*293-derived fluorescent strains expressing YFP (TBK1.1) and RFP (TBK5.1) that mimic pathogenesis of the parental strain in larval zebrafish ^8^, as well as the auxotrophic *Af*293.1 *pyrG-*strain ^43^. Spores were prepared by spreading 10^6^ spores onto 10 cm glucose minimal media (GMM) plates (with uracil and uridine added to grow pyrG-spores) and incubating at 37°C for 3-4 days. The spores were then collected into sterile water with 0.01% Tween, filtered through miracloth, centrifuged, washed with sterile PBS, and resuspended in PBS to a final concentration of 1.5 × 10^8^ spores/ml and stored for up to ∼4 weeks, as previously described ^31^.

**Table 5:** *Aspergillus fumigatus* strains used in this study.

| Parental background | Strains | Genetic background | Fluorescence | References |
| --- | --- | --- | --- | --- |
| *Af293* | *Af293.1* | *pyrG1* | None | 43 |
| *Af293* | TBK1.1 | *pyrG1, gpdA::YFP::A. fumigatus pyrG* | YFP | 30 |
| *Af293* | TBK5.1 | *pyrG1, argB1, ΔakuA, A. fumigatus argB, gpdA::mCherry::A. fumigatus pyrG* | mCherry | 8 |

### Conjugation of AlexaFluor633 to spores

Spores from an *Af293*-derived fluorescent strain expressing YFP (TBK1.1) were coated with AlexaFluor633 as described previously ^44^. Spores and biotin-XX, SSE (Molecular Probes) were incubated on a shaker in 0.05 M NaHCO_3_ at 4°C for 1.5–2 hours. The spores were pelleted and washed twice with 100 mM Tris-HCl (pH 8.0) on a shaker at 4°C for 30-45 min to remove unbound biotin. The spores were washed with 1x PBS twice and resuspended in 1x PBS containing 20 µg/ml of streptavidin-AlexaFluor633 (Invitrogen) and subsequently incubated on a shaker for 40 min at room temperature. The labeling of spores was confirmed by fluorescence microscopy. The stained spores were then pelleted, washed and resuspended in 1x PBS, and a spore suspension was made and stored as 1.5 × 10^8^ spores/ml at 4°C for up to ∼4 weeks.

### Zebrafish larval hindbrain microinjection

2-day post-fertilization (dpf) zebrafish larvae were anesthetized with tricaine and injected with *A. fumigatus* spores as previously described ^31^. Spore suspensions at a concentration of 1.5 × 10^8^ spores/ml were mixed 2:1 with 1% phenol red to achieve a final concentration of 1 × 10^8^ spores/ml. Injection plates were prepared with 2% agarose in E3 medium, coated with 2% bovine serum albumin (BSA), and rinsed. Anesthetized larvae, aligned laterally on an agarose plate, were injected in the hindbrain ventricle with spores (targeting ∼50–100 spores per larva), and the actual doses were monitored by randomly selecting 8 larvae for CFU plating as described below immediately after injection and these CFUs are reported in the figure legends. For mock infections, PBS mixed with phenol red was used. Post-injection, larvae were rinsed twice with E3 medium without methylene blue to remove tricaine and any residual spores. Injected larvae were maintained in 96-well plates for survival and CFU assays, and in 35-mm dishes or 48-well plates for imaging analyses.

### Bafilomycin A1 treatment

Bafilomycin A1 (BafA1) (Sigma-Aldrich), was resuspended in DMSO at 50 µM or 100 µM, stored at -20°C, and diluted into E3 without methylene blue to concentrations of 50 nM or 100 nM (0.1% DMSO). Larvae were then placed in E3 with diluted drug 1 hour post injection (hpi). DMSO at 0.1% was used as a vehicle control.

### CFU analysis

To monitor colony-forming units (CFUs), larvae were placed singly in 1.7 mL microcentrifuge tubes containing 90 µL PBS with 1 mg/mL ampicillin and 0.5 mg/mL kanamycin and homogenized in a TissueLyser (Qiagen) at 1,800 oscillations/min (30 Hz) for 12 min. Samples were centrifuged at 17,000 g for 30-45 seconds, resuspended and spread on glucose minimal media (GMM) plates and incubated at 37°C for 3 days before counting. For all survival analyses, CFU counts were done on eight larvae from each injection condition immediately after injection to determine the actual spore dose, and the average dose is reported in each figure legend. To analyze spore killing ability of macrophages, *Af*293.1 *pyrG-* spores which do not germinate well in vivo were used ^8^. From 0 to 5 dpi, twelve larvae were plated per day, condition, and replicate on GMM plates supplemented with uridine and uracil. The number of fungal colonies was counted and normalized to the average CFU burden at 0 dpi to calculate the percent initial spore burden.

### Tail wounding

Tail transection was performed on anesthetized 2 dpf larvae with a no.10 feather surgical blade under a dissection light microscope. Larvae were then transferred to fresh E3 medium, kept at 28°C, and fixed 3 hours post-wounding (hpw) with 4% formaldehyde in 1× PBS for 1 hour on a rocker at room temperature. Larvae were then washed 3 times in 1× PBS for 5 min on a rocker at room temperature and immediately used for confocal imaging or stored in PBS-Tween at 4°C until imaging.

### TUNEL staining

Terminal deoxynucleotidyl transferase dUTP Nick-End Labeling (TUNEL) staining for whole larvae was adapted and modified from previous studies ^45,46^. At 2 dpf, larvae were anesthetized and fixed overnight in 4% paraformaldehyde (PFA; Polysciences, Inc) in PBST (0.05 % Tween20) at room temperature on a rocker. Larvae were then dehydrated in ascending grades of 25%, 50%, 75%, and 100% methanol for 5 min each. Larvae were rehydrated through the same descending grades of methanol back to PBST followed by permeabilization in 0.1 % sodium citrate (pH 6.0)(Fisher BioReagents) for 15 min at room temperature. The larvae were digested with Proteinase K (1 µg/mL in PBST)(NEB) for 15 min. After three PBST washes, TUNEL labeling was performed using the In Situ Cell Death Detection Kit, TMR Red (Roche) with 45 µL Label solution and 5 µL TdT enzyme per sample to label DNA strand breaks followed by incubation for 1 hour at 37°C in the dark. Larvae were washed three times in PBST prior to imaging.

### Confocal microscopy

For all imaging experiments, larvae were treated with 200 µM N-phenylthiourea (PTU) from 24 hours post-fertilization to inhibit pigmentation. All imaging was performed on a Zeiss Cell Observer Spinning Disk confocal microscope equipped with a Photometrics Evolve 512 EMCCD camera to acquire multi-channel Z-stack images. For time-lapse imaging, larvae were anesthetized and mounted in glass-bottom dishes in 1% low-melting agarose in E3 with PTU and tricaine. Larvae were imaged with a Plan-Apochromat 10x objective (0.3 NA) for 12 to 18 hours at 4-5 min intervals. For single time point or daily live imaging of the infection site, or for imaging of fixed larvae, larvae were individually anesthetized in E3 medium containing tricaine and placed in chambers of a glass-bottom zWEDGI device ^47^ and imaged with a Plan-Apochromat 20x objective (0.8 NA) or EC Plan-Neofluar 40x objective (0.75 NA). For imaging of macrophage-spore interactions at 1 dpi, larvae were stained with 10 µM Lysotracker Red DND-99 (Invitrogen) diluted into E3 at 28°C for 1 hour and then rinsed twice in E3 prior to anesthesia and imaging. For multi-day live imaging, larvae were removed from the zWEDGI device after imaging, washed in E3 without methylene blue, and returned to the same well with E3 containing PTU and placed back at 28°C. For imaging of the whole body, larvae were also placed in a zWEDGI device a Plan-Apochromat 10x objective (0.3 NA) was used and 4-5 images spanning the body were acquired.

### Image analysis

Image analysis was conducted using ImageJ/Fiji software (ImageJ version 1.54i with Java version 1.8.0_172) ^48^. Macrophage recruitment to the infection site in the hindbrain ventricle was quantified by manually counting fluorescent-positive cells or by fluorescence intensity thresholding to measure macrophage area within the ROI of the hindbrain. Interaction of macrophages with spores was manually quantified as fluorescent-positive macrophages proximal to fluorescent-positive spores during 1-hour windows. The percentage of spores internalized by macrophages, the percentage of alive versus dead spores, and the percentage of spores in acidified Lysotracker+ compartments were all also determined by manual counting. Fungal burden was quantified as fluorescent-positive area by intensity thresholding within ROIs containing hyphae and by manual classification of the stage of fungal development and growth within larvae. If a single spore within a larva progressed to a new developmental stage, we classified the entire larva as having reached that stage. For quantification of macrophages and neutrophils across the whole larvae, multiple images were pieced together to span the entire body and fluorescent-positive cells were manually counted. For analysis of TUNEL staining, overlap of TUNEL signal and H2B-GFP signal was manually counted and total TUNEL+ area was quantified by intensity thresholding within an ROI of the caudal hematopoietic tissue (CHT). Macrophage recruitment to tail wounds was also quantified by manual counting of fluorescent-positive cells. For some analyses and for display of images, maximum intensity Z-projections were generated, and images were resized to 1024 x 1024 pixels via bilinear interpolation.

### Graphs and statistical analysis

Statistical analyses were conducted on data pooled from three independent replicates and all graphs present pooled data from all replicates. Data was analyzed and graph generation was performed using R studio (version 2024.04.0 Build 735; Chocolate Cosmos) with R version 3.5.2. Survival data and survival curves were analyzed by Cox proportional hazard regression as previously described regression as previously described ^8,31^. In these analyses, experimental replicate was included as a grouping variable, with either *atp6v1h* and *luc* larvae as the two experimental groups, or larva treated with two concentrations of Baf A1 or PBS control, and hazard ratios were calculated to represent the relative instantaneous risk of death between conditions. The total number of larvae across all three replicates for each survival experiment (n) is indicated in each figure.

For analysis of imaging and CFU data, estimated marginal means (emmeans) and standard error of the mean (SEM) were calculated for all experimental conditions, and pairwise comparisons were performed by ANOVA using Tukey’s adjustment. Fisher’s exact test was used to assess differences in outcomes between *irf8* genotypes. For single time point imaging experiments, the data each individual larvae are presented from three replicates, color coded by replicate. Statistical significance is represented by asterisk marks (*p<0.05, **p<0.01,***p<0.001,****p<0.0001) in the figure and statistical values written in figure legends.

## Results

### v-ATPase complex colocalizes with *A. fumigatus* spores in macrophages in larval zebrafish

Macrophages are the first innate immune cells to respond to *A. fumigatus* infection, playing a crucial role in phagocytosing spores and inhibiting their germination ^8,27,50^. One of the major mechanisms that macrophages use to control phagocytosed targets is phagosomal acidification, mediated by localization of the v-ATPase complex to the phagosome. A proteomic analysis found components of the v-ATPase complex in isolated phagosomes from an *A. fumigatus*-infected murine macrophage cell line ^51^, however, this localization has not been confirmed in a whole animal model. To visualize v-ATPase localization to phagosomes containing *A. fumigatus* spores in vivo, we generated a new transgenic zebrafish line to express Atp6v1h tagged with TagRFPT under a macrophage specific promoter (*mfap4:TagRFPT.zf1-atp6v1h*). Atp6v1h encodes the regulatory H subunit of the V1 domain of the v-ATPase complex, which is essential for ATP hydrolysis ^52^. We then infected larvae with spores of a YFP-expressing *Af*293-derived strain of *A. fumigatus* at 2 days post-fertilization (dpf) via hindbrain injection and performed confocal imaging and quantitative analysis of TagRFPT-Atp6v1h co-localization with *Aspergillus* spores in macrophages from 1 to 5 days post-infection (dpi). As previously reported, macrophages migrate to the infection site, phagocytose spores, and can form small clusters by 1 dpi (Fig 1A) ^8^. In these clusters, spores with TagRFPT-Atp6v1h co-localization and those without co-localization can both be found (Fig 1A). Also consistent with previous studies, spores persist for up to 5 dpi, with the total number of spores per larvae slowly decreasing as some are cleared over time (Fig 1B) ^8^. However, the percentage of spores in macrophages that co-localize with TagRFPT-Atp6v1h is relatively consistent across the 5 days of imaging, ranging from ∼35-50% of spores (Fig 1C). These results suggest that the v-ATPase complex is persistently recruited to macrophage phagosomes containing *Aspergillus* spores, as macrophages slowly clear spores, supporting the idea that this complex plays a role in macrophage control of infection.

**Figure 1:**
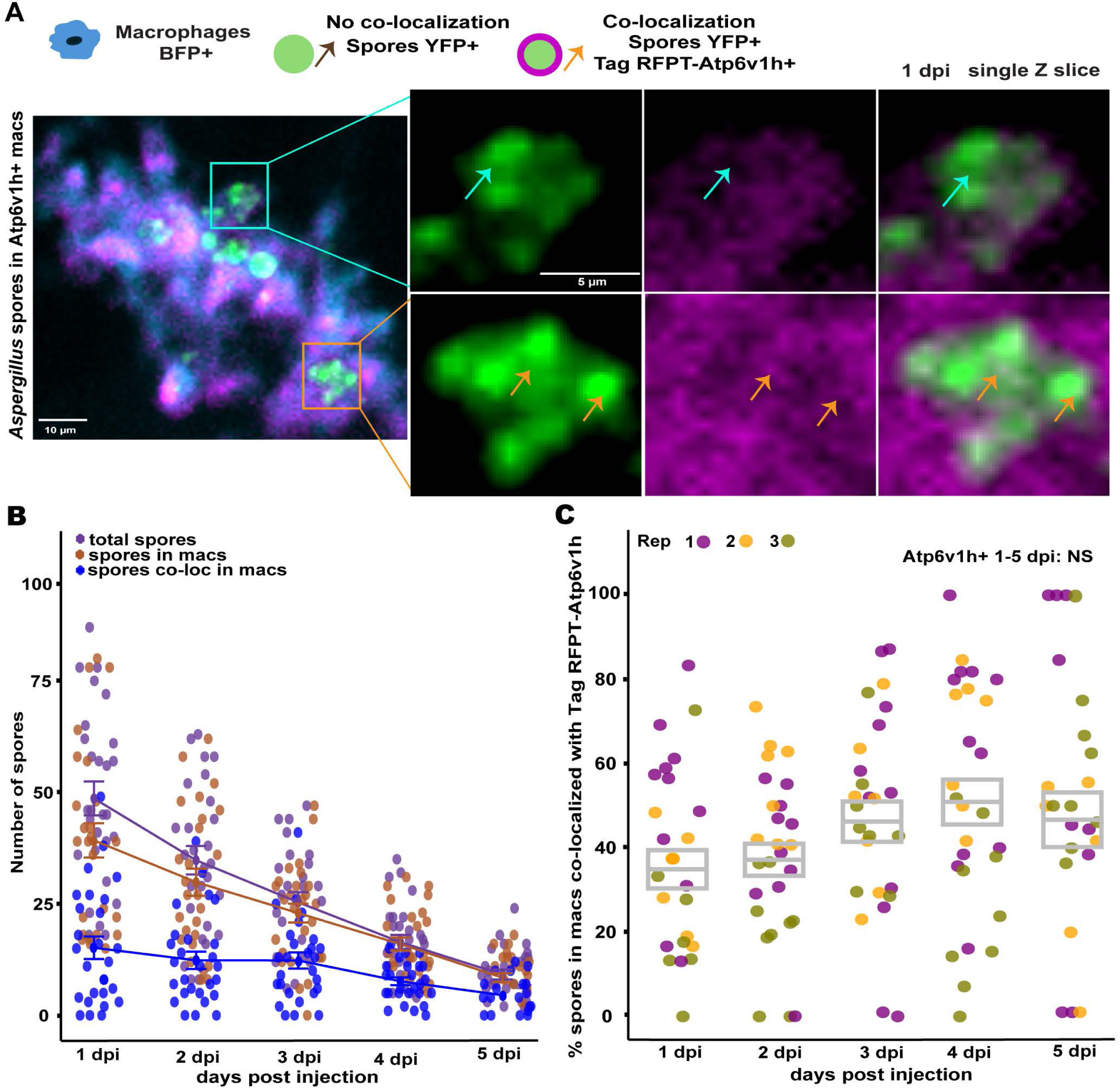
Fluorescently-tagged Atp6v1h co-localizes with spores in macrophages across 5 days of infection. Transgenic larvae expressing TagRFPT-Atp6v1h (*mfap4:TagRFPT-atp6v1h*) and with fluorescently-labeled macrophages (*mfap4:BFP*) were injected with *Af*293-derived YFP-expressing *A. fumigatus* spores (TBK1.1) and imaged from 1 to 5 dpi. (A) Representative single z-slice confocal images from 1 dpi. Orange arrows indicate spores that co-localized with TagRFPT-Atp6v1h and cyan arrows indicate spores without co-localization. Left scale bar: 10 µm; right scale bar: 5 µm. (B) Quantification of total spores (purple), spores within macrophages (brown), and TagRFPT-Atp6v1h-co-localized spores within macrophages (blue). Each dot represents the spore count from an individual zebrafish larva. Lines and error bars indicate the emmeans ± SEM for each group at each time point. (C) Quantification of the percentage of spores inside of macrophages that co-localize with TagRFPT-Atp6v1h. Each dot represents an individual larva, with colors indicating separate experimental replicates. *P* values were calculated by emmeans and ANOVA, gray boxes represent the emmeans ± SEM.

### v-ATPase deficiency impairs macrophage-mediated control of *A. fumigatus* infection

In vitro studies have demonstrated that inhibiting v-ATPase activity of macrophages with bafilomycin A1 (Baf A1) reduces their spore killing ability ^53–55^. However, macrophages in zebrafish have lower spore killing ability (Fig 1B) ^8^, and the role of the v-ATPase complex in macrophage-mediated defense in a whole animal is also unclear. To investigate these questions, we targeted *atp6v1h* using CRISPR-Cas9 by injecting single-celled embryos with Cas9 protein and two gRNAs (Supp Fig 1A, B). PCR, gel electrophoresis, and sequence analysis of the DNA sequences targeted by these gRNAs confirmed high efficiency targeting of this gene, with an average editing efficiency of ∼57% for gRNA1 and ∼69% for gRNA2 (Suppl Fig 1C, D). The larvae resulting from this embryo injection are therefore termed *atp6v1h* crispant larvae. gRNAs targeting the *luciferase* coding sequence were used as a control. We first generated *atp6v1h* and control *luc* crispants in wild-type larvae, infected these larvae with spores of an *Af*293-derived strain of *A. fumigatus* at 2 days post-fertilization (dpf) via hindbrain injection, and monitored survival for 7 days. Control wild-type larvae are largely resistant to *A. fumigatus* infection, with a survival rate of 96% (Fig 2A). Targeting *atp6v1h* significantly reduced survival to 82%, and *atp6v1h* crispants are 6.4 times more likely to succumb to infection compared to *luc* control larvae (Fig 2A).

**Figure 2:**
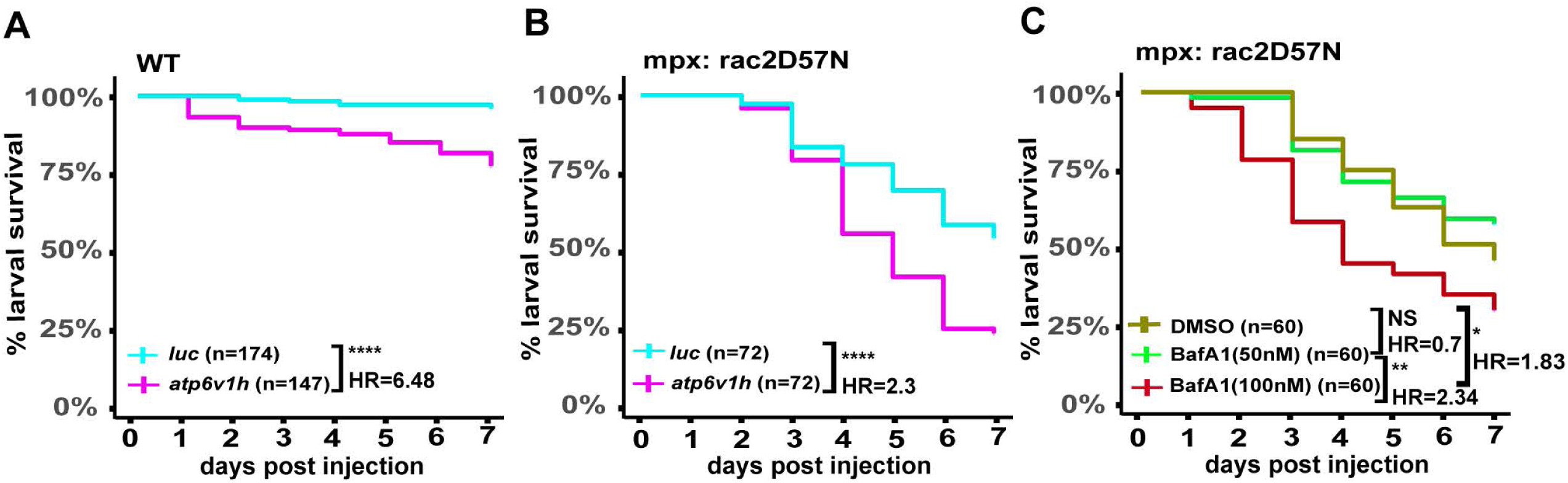
v-ATPase deficiency increases larval susceptibility to *A. fumigatus* infection. (A) *atp6v1h* crispants or *luc* controls in a wild-type background were infected with *Af*293-derived *A. fumigatus* spores (TBK1.1) and monitored for survival for 7 days post injection (dpi). Average injected CFUs: *atp6v1h* = 92; *luc* = 89. (B) *atp6v1h* crispants or *luc* controls in a neutrophil-defective background (*mpx:rac2D57N*) were infected with *Af*293-derived *A. fumigatus* spores (TBK1.1) and monitored for survival for 7 dpi. Average injected CFUs: *atp6v1h* = 73; *luc* = 74. (C) Larvae in a neutrophil-defective background (*mpx:rac2D57N*) were infected with *Af*293-derived *A. fumigatus* spores (TBK1.1), treated with Bafilomycin A1 (BafA1) or DMSO vehicle control, and monitored for survival for 7 dpi. Average injected CFUs = 80. *P* values and hazard ratios were calculated by Cox proportional hazard analysis. Each line represents the combined data from three independent replicates.

To investigate the impact of v-ATPase dysfunction on macrophage-specific responses to *A. fumigatus* infection, we next generated *atp6v1h* crispants in a neutrophil-defective zebrafish line (*mpx:mCherry-2A-rac2^D57N^*). While wild-type larvae have intact macrophage and neutrophil function, the *rac2^D57N^*line relies solely on macrophages due to impaired neutrophil migration caused by a dominant-negative Rac2 variant expressed specifically in neutrophils ^38^. Following infection of *atp6v1h* crispants and *luc* control *rac2^D57N^* larvae with *A. fumigatus* spores, only 25% of *atp6v1h* crispants survived, whereas the *luc* control group exhibited a 55% survival rate, indicating a significant 30% decrease in survival associated with *atp6v1h* deficiency (Fig 2B). Neutrophil-defective larvae that lack v-ATPase function are 2.3 times more likely to succumb to infection compared to control *rac2^D57N^* larvae, highlighting the critical role of v-ATPase function in macrophage-mediated defense against *A. fumigatus*.

To confirm the role of the v-ATPase complex in controlling *A. fumigatus* infection, we pharmacologically inhibited v-ATPase function with the macrolide antibiotic bafilomycin A1 (BafA1) ^56,57^. Larvae were exposed to concentrations of BafA1 that are not toxic to fish and are similar to effective doses in vitro ^49,58^. In neutrophil-defective *rac2^D57N^* larvae infected with *A. fumigatus* spores, 100 nM BafA1 significantly increased larval susceptibility, with mortality rates 20-30% higher than those treated with 50 nM BafA1 or DMSO vehicle control (Fig 2C). BafA1 treatment did not significantly affect survival of larvae mock-infected with PBS (Suppl. Fig 2). These data confirm that inhibition of v-ATPase function compromises fungal control in neutrophil-defective larvae that rely on macrophage-mediated defense against *A. fumigatus*.

### v-ATPase deficiency impairs macrophage recruitment and spore interaction in early immune response

We next wanted to understand how macrophage function is compromised in v-ATPase deficient *A. fumigatus-*infected larvae. We first utilized live imaging to monitor the initial recruitment and behavior of macrophages in *atp6v1h* crispants and *luc* control larvae from 1-18 hours post injection (hpi). Larvae with fluorescently-labeled macrophages (*mpeg1:H2B-GFP*; *mfap4:BFP*) were injected with RFP-expressing *Af*293-derived *A. fumigatus* spores (Fig 3A). Starting at 6 hpi, control *luc* larvae have significantly higher numbers of macrophages at the infection site compared to *atp6v1h* crispants (Fig 3B; Video 1A, 1B). This significant difference in macrophage recruitment persisted until the end of the experiment at 18 hpi (Fig 3B). In control *luc* larvae, recruited macrophages form clusters (Fig 3A, Video 1A) while in *atp6v1h* crispants, macrophages remain separate and do not form such clusters (Fig 3A, Video 1B). As we observed significantly fewer macrophages at the infection site with altered clustering behavior in *atp6v1h* crispants, we next determined if fewer macrophages interact with *A. fumigatus* spores in *atp6v1h* crispants. In one hour windows across the timelapse, we quantified the number of macrophages around spores. In *luc* control larvae, the number of interacting macrophages steadily increases from ∼5 interacting macrophages at 1 hpi to ∼20 interacting macrophages at 18 hpi (Fig 3C). The number of macrophages interacting with spores was significantly reduced in *atp6v1h* crispant larvae, with fewer than ∼10 interacting macrophages at all time points (Fig 3C). These findings suggest that v-ATPase deficiency impairs both macrophage numbers in the infected hindbrain and macrophage interactions with fungal spores.

**Figure 3:**
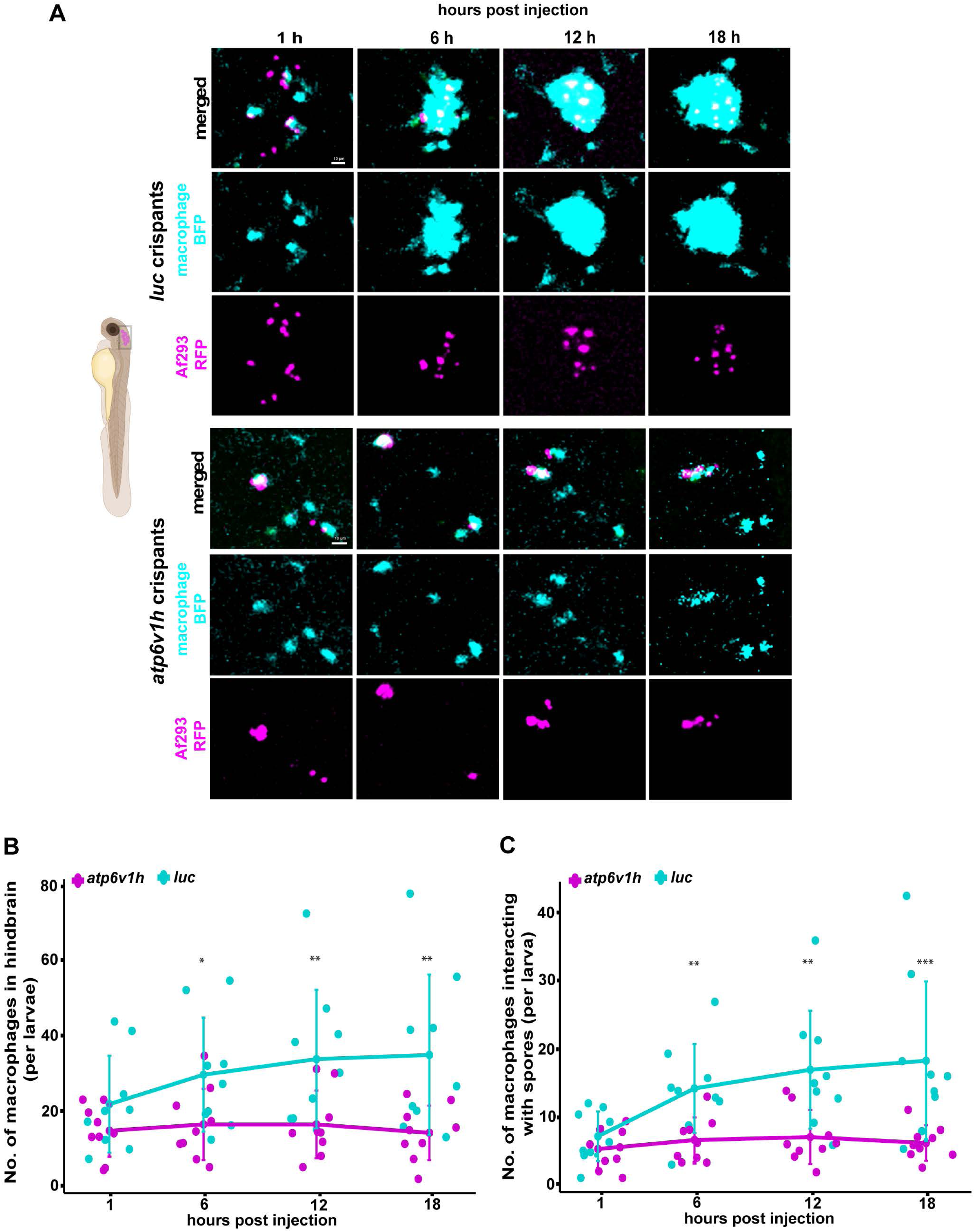
v-ATPase deficiency impairs early macrophage recruitment and interaction with spores. *atp6v1h* crispants or *luc* control larvae with fluorescently-labeled macrophages (*mfap4:BFP*; *mpeg1:H2B-GFP*) were infected with RFP-expressing *Af*293-derived *A. fumigatus* spores (TBK5.1) and followed by confocal live timelapse imaging from 1-18 hours post injection. (A) Representative confocal images of BFP+ macrophages recruited to RFP-expressing spores in the hindbrain. Scale bar: 10 µm. (B, C) Macrophage number and macrophage-spore interactions in the hindbrain of larvae were manually quantified across one-hour windows centered at 1, 6, 12, and 18 hours post injection. Each dot represents an individual larva, with colors indicating separate experimental replicates. *P* values were calculated by emmeans and ANOVA, bars represent the emmeans ± SEM. Asterisks (*) denote a statistically significant difference between *luc* controls and *atp6v1h* crispants at a given time point.

### Larvae deficient in v-ATPase have lowered overall macrophage and neutrophil numbers

As we observed decreased macrophage recruitment to *A. fumigatus* spores in *atp6v1h* crispants, we wondered if this was due to a specific defect in migration or if v-ATPase deficiency affects overall macrophage numbers. To quantify total macrophage numbers, we generated *atp6v1h* crispants and *luc* controls in a transgenic line with fluorescently-labeled macrophages (*mpeg1:GFP-H2B*), imaged larvae at 2 dpf, and counted macrophages across the whole body (Fig 4A). The total number of macrophages per larvae was significantly reduced in *atp6v1h* crispants compared to *luc* control larvae (Fig 4B). On average, *atp6v1h* crispants had approximately 70 fewer macrophages per larvae, with average counts of 153 in *atp6v1h* crispants compared to 224 in *luc* controls (Fig 4B). Assessment of neutrophil numbers in a fluorescent neutrophil transgenic line (*lyz:H2B-mCherry*) also revealed defects in neutrophil numbers in *atp6v1h* crispant larvae compared to *luc* controls, however this effect was not as stark (Suppl Fig 3).

**Figure 4:**
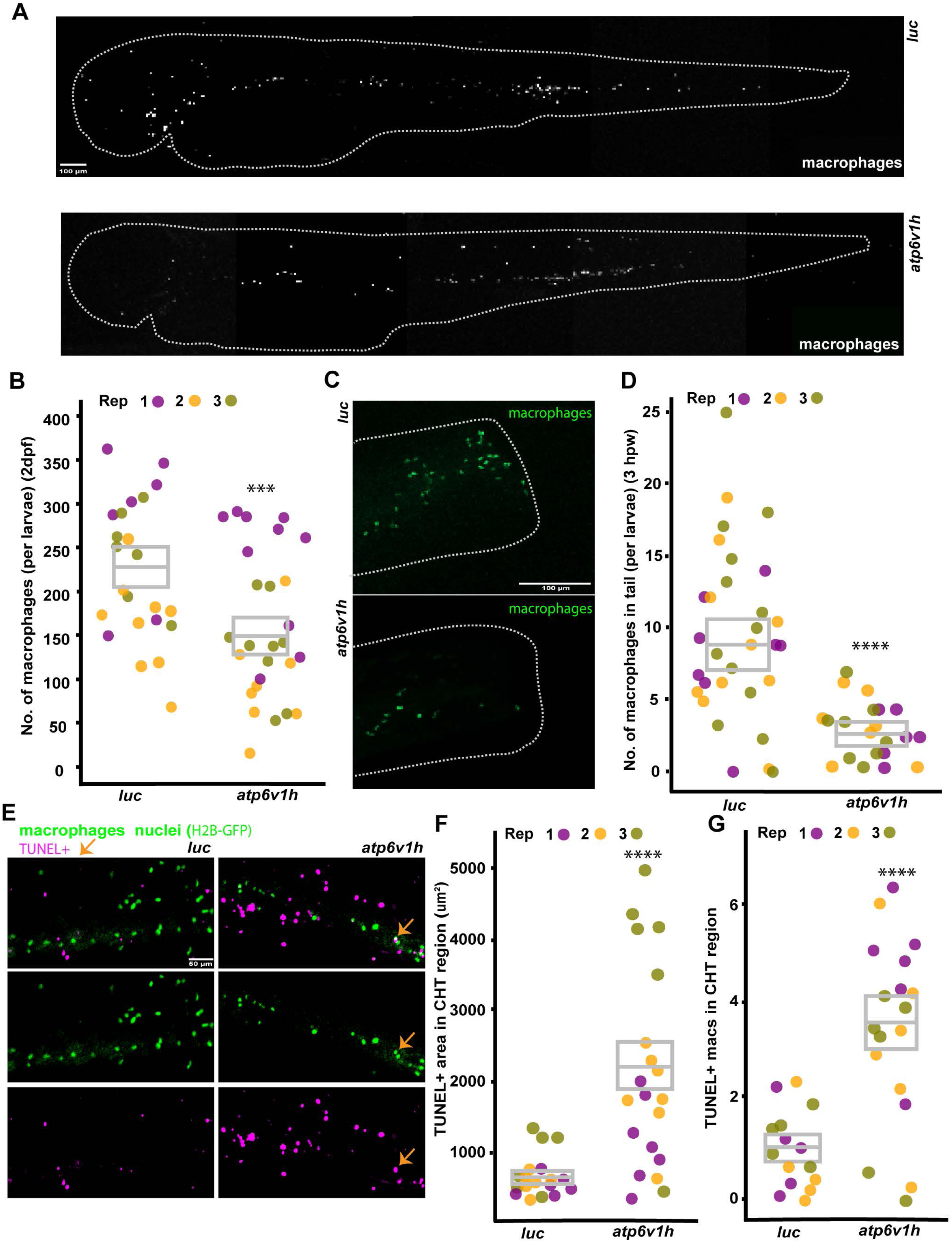
v-ATPase deficiency impairs overall macrophage numbers and response to injury. (A, B) *atp6v1h* crispants and *luc* controls with fluorescently-labeled macrophages (*mpeg1:H2B-GFP*) were imaged at 2 days post-fertilization (dpf). Representative confocal images are shown,, scale bar: 100 µm (A), and total macrophage numbers per larva were quantified (B). Each dot represents an individual larva, with colors indicating separate experimental replicates. *P* values were calculated by emmeans and ANOVA, and gray boxes represent emmeans ± SEM. (C, D) At 2 dpf, *atp6v1h* crispants and *luc* control larvae, each expressing *mpeg1:H2B-GFP* labeled macrophages, underwent tail transection, and the wound site was imaged 3 hours post-wounding. Representative confocal images are shown, scale bar: 100 µm (C) and macrophage numbers recruited to tail wound per larva were quantified (D). Each dot represents an individual larva, with colors indicating separate experimental replicates. *P* values were calculated by emmeans and ANOVA, and gray boxes represent emmeans ± SEM. (E-G) At 2 dpf, *atp6v1h* crispants and *luc* control larvae, each expressing *mpeg1:H2B-GFP* labeled macrophages, were used for TUNEL staining. Representative confocal images in the caudal hematopoietic tissue (CHT) are shown, scale bar: 50 µm (E) and total TUNEL+ positive area in the CHT (F) and number of TUNEL+ macrophages in the CHT (G) were quantified. Each dot represents an individual larva, with colors indicating separate experimental replicates. *P* values were calculated by emmeans and ANOVA, gray boxes represent the emmeans ± SEM.

We next determined whether the reduced macrophage recruitment to *A. fumigatus* spore infection that we observed (Fig 3) was specific to infection or due to a broader defect in macrophage numbers and migration. We therefore assessed macrophage recruitment to a tail wound in both *atp6v1h* crispants and *luc* control larvae in a transgenic line with fluorescently-labeled macrophages (*mpeg1:GFP-H2B*). At 3 hours post wounding (hpw), the number of macrophages recruited to the injury site was significantly reduced in *atp6v1h* crispant larvae compared to *luc* controls (Fig 4C, D). On average, *luc* control larvae recruited ∼8 macrophages to the tail wound, whereas *atp6v1h* crispants recruited only ∼3-4 macrophages to the tail wound (Fig 4D). These data suggest that v-ATPase function is essential for general macrophage responses to injury, likely due to defects in total cell numbers in larvae.

To determine if total cell numbers are decreased due to increased macrophage apoptosis, we performed TUNEL staining on 2 dpf larvae with fluorescently-labeled macrophages (*mpeg1:GFP-H2B*) and measured staining in the caudal hematopoietic tissue (CHT), the site of macrophage development (Fig 4E). The total TUNEL+ area within the CHT region was significantly increased in *atp6v1h* crispants compared to *luc* controls (Fig 4F). Focusing on macrophages, the number of TUNEL+ nuclei increased from ∼1 per larvae in *luc* controls to ∼3.5 in *atp6v1h* crispants (Fig 4G). These findings demonstrate that *atp6v1h* deficiency does increase overall apoptosis in the CHT, but the increase in macrophage apoptosis likely does not fully account for the overall difference in macrophage numbers (Fig 4B). Therefore, macrophage development is also likely compromised in *atp6v1h* crispant larvae.

### v-ATPase function is required for acidification but not killing of macrophage-internalized spores

While v-ATPase impairment reduces overall macrophage numbers and macrophage recruitment to the infection site, we still sought to determine the effect of v-ATPase deficiency on the function of macrophages that successfully reach the infection site. We therefore focused on macrophages at the infection site at 1 dpi, again in neutrophil-defective *rac2^D57N^* larvae in which macrophages are the only major immune cell present. Macrophages were again fluorescently-labeled (*mfap4:BFP*), and to track the outcome of spores, we used live-dead staining with *Af*293-derived *A. fumigatus* spores expressing YFP and with a cell wall label of AlexaFluor633 (Fig 5A), as previously described^8,44^. Additionally, we assessed lysosomal acidification by staining larvae with Lysotracker Red prior to imaging (Fig 5A). In line with our previous findings from timelapse microscopy, control *luc* larvae exhibited significantly greater macrophage recruitment at the infection site compared to *atp6v1h* crispants (Fig 5B) and a significantly higher percentage of spores was internalized by macrophages in *luc* controls compared to *atp6v1h* crispants (Fig 5C). Therefore, to specifically investigate how v-ATPase deficiency affects the processing of *A. fumigatus* spores by macrophages, we focused exclusively on the spores that were inside of macrophages. Of spores found in macrophages, ∼40% were found in acidified compartments in *luc* control larvae (Fig 5A, D). However, only ∼25% of spores were acidified in *atp6v1h* crispant larvae, a significant decrease (Fig 5A, D), demonstrating impaired phagolysosomal acidification of spores in v-ATPase-deficient macrophages. As the v-ATPase complex is the main driver of lysosomal acidification, we also sought to assess the general effect of *atp6v1h* CRISPR-targeting on lysosomal acidification in macrophages. Total lysotracker-positive area in macrophages was significantly reduced in *atp6v1h* crispants compared to *luc* controls, both in absolute terms and when normalized to total macrophage area (Suppl Fig 4A, B) demonstrating that *atp6v1h* crispants have overall impaired v-ATPase function.

**Figure 5:**
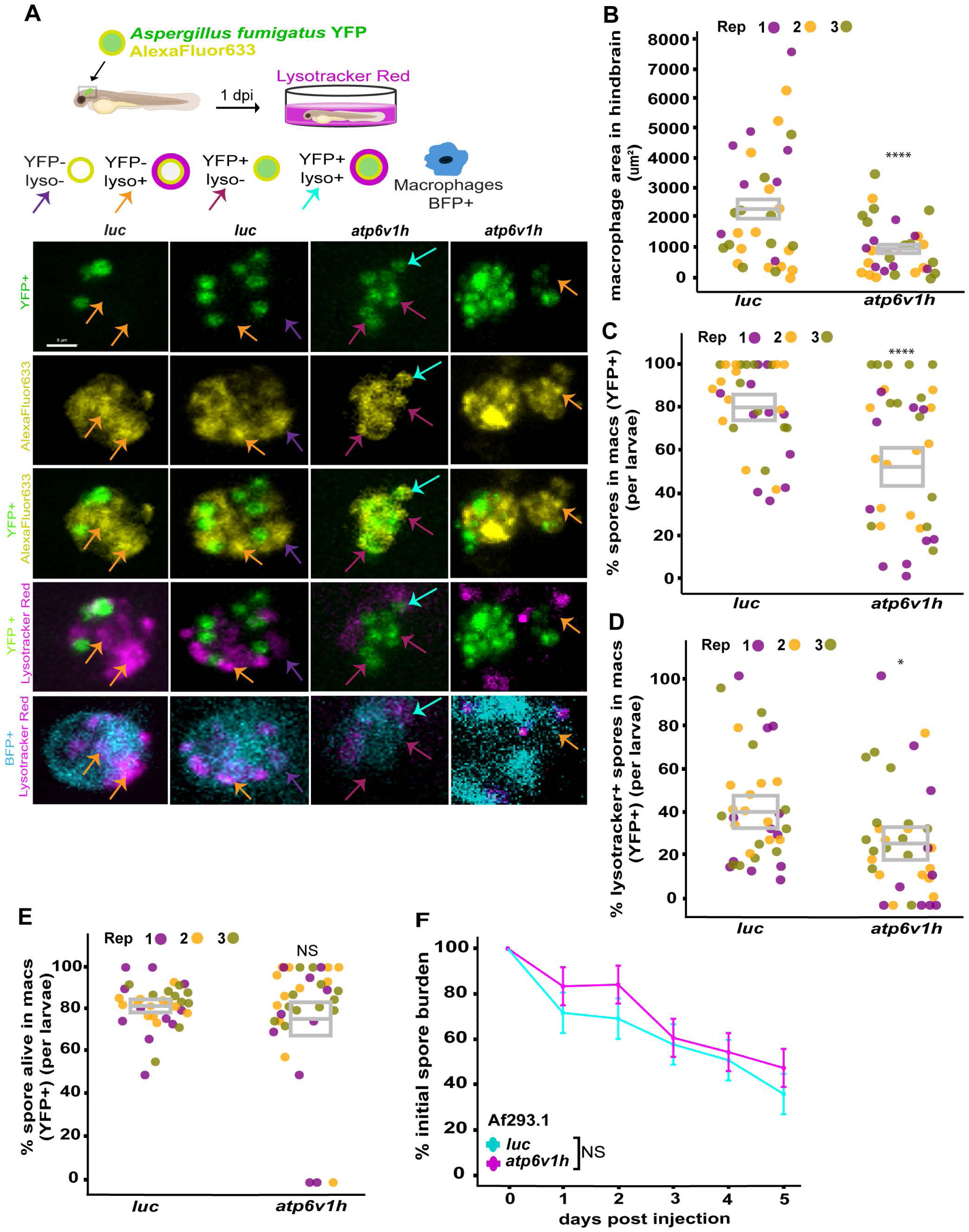
v-ATPase function promotes macrophage-mediated spore acidification but not spore killing. (A-E) *atp6v1h* crispants and *luc* controls with fluorescently-labeled macrophages (*mfap4:BFP*) were injected with *Af*293-derived YFP-expressing *A. fumigatus* spores (TBK1.1) with AlexaFluor633 conjugated to the cell wall. At 1 dpi, larvae were stained with Lysotracker Red dye for 1 hour and then confocal imaged. Experimental setup and representative images are shown (A). Different color arrows indicate spores in different categories based on fluorescence. Scale bar: 5 µm. Macrophage area in the hindbrain region (B), percentage of spores in the hindbrain (both YFP+ and YFP-) that were internalized by macrophages (C), percentage of macrophage-internalized spores that are lysotracker+ (D), and percentage of spores inside macrophages that are alive (E) were quantified. Each dot represents an individual larva, with colors indicating separate experimental replicates. *P* values were calculated by emmeans and ANOVA, and bars represent the emmeans ± SEM. (F) *atp6v1h* crispants and *luc* controls in a wild-type background were infected with *Af*293-derived *A. fumigatus Af*293.1 *pyrG-* spores and spore killing was monitored by plating CFUs from 12 individually homogenized larvae each day for 5 dpi. Average injected CFUs: *atp6v1h* = 90; *luc* = 83. *P* values were calculated by emmeans and ANOVA and emmeans ± SEM are plotted.

Surprisingly, despite reduced lysosomal acidification of spores in *atp6v1h* crispants, we found no significant change in spore killing by macrophages between *atpv1h* crispants and *luc* controls (Fig 5E). These data suggest that while v-ATPase function promotes spore acidification, it is not required for macrophage-mediated spore killing. To confirm that v-ATPase complex function is not required for macrophage-mediated spore killing, we quantified colony forming units (CFUs) from homogenized single larvae over a 5-day infection period. To specifically focus on spore killing, we infected larvae with an *Af*293-derived pyrG-strain, which is auxotrophic for uracil and uridine and does not efficiently germinate in the larval zebrafish model ^8^. Again, we observed no significant defect in spore clearance in *atp6v1h* crispants compared to *luc* controls (Fig 5F). Both *atp6v1h* crispants and *luc* controls exhibited a gradual decline in spore burden over five days post injection (Fig 5F). Overall, these findings demonstrate that while v-ATPase deficiency disrupts macrophage recruitment, spore engulfment, and lysosomal acidification, it does not significantly impact spore killing, suggesting that acidification alone may not be the primary determinant of spore killing within macrophages.

### v-ATPase function promotes macrophage control of fungal germination and development

As *atp6v1h* deficiency impacts spore uptake and acidification without impairing spore killing within macrophages, we wondered what the cause of reduced larval survival in *atp6v1h* crispants is. We hypothesized that neutrophil-defective *atp6v1h* crispants have greater spore germination and hyphal development leading to larval death. To test this hypothesis, we generated *atp6v1h* crispants and *luc* controls in neutrophil-deficient larvae with fluorescently-labeled macrophages (*mfap4:BFP*), infected larvae with *Af*293-derived *A. fumigatus* spores expressing YFP and performed daily live imaging for five consecutive days to track fungal development and growth (Fig 6A). Fungal development was classified as dormant spores, swollen spores, germinating spores, invasive hyphal growth, and larval death, with each larva assigned a category based on the most advanced stage reached by any fungal unit with the larva. In comparison to *luc* controls, *atp6v1h* crispants experienced more rapid infection progression (Fig 6A, B). At 1 dpi, 90% of *luc* control larvae only had dormant spores, with 10% having swollen spores at the infection site. In contrast, only 50% of *atp6v1h* crispants still only had dormant spores, with 10% having swollen spores, 30% with germinating spores, 3% with invasive hyphae, and 12% already having succumbed to the infection, indicating accelerated infection progression in *atp6v1h* crispants. At 2 dpi, 30% of *atp6v1h* crispants were already experiencing invasive hyphae, while this stage of development was only present in <20% of *luc* control larvae. By 4 dpi, no spores in *atp6v1h* crispants remained in the dormant stage, whereas some *luc* control larvae retained dormant spores up to 5 dpi (Fig 6B). In these larvae, we also quantified fungal burden by measuring the fluorescent-positive fungal area (Fig 6C). These measurements show a similar trend, with *atp6v1h* crispants experiencing accelerated fungal growth which peaks at 2–3 dpi, whereas *luc* controls experience this growth later, at 3–5 dpi (Fig 6C).

**Figure 6:**
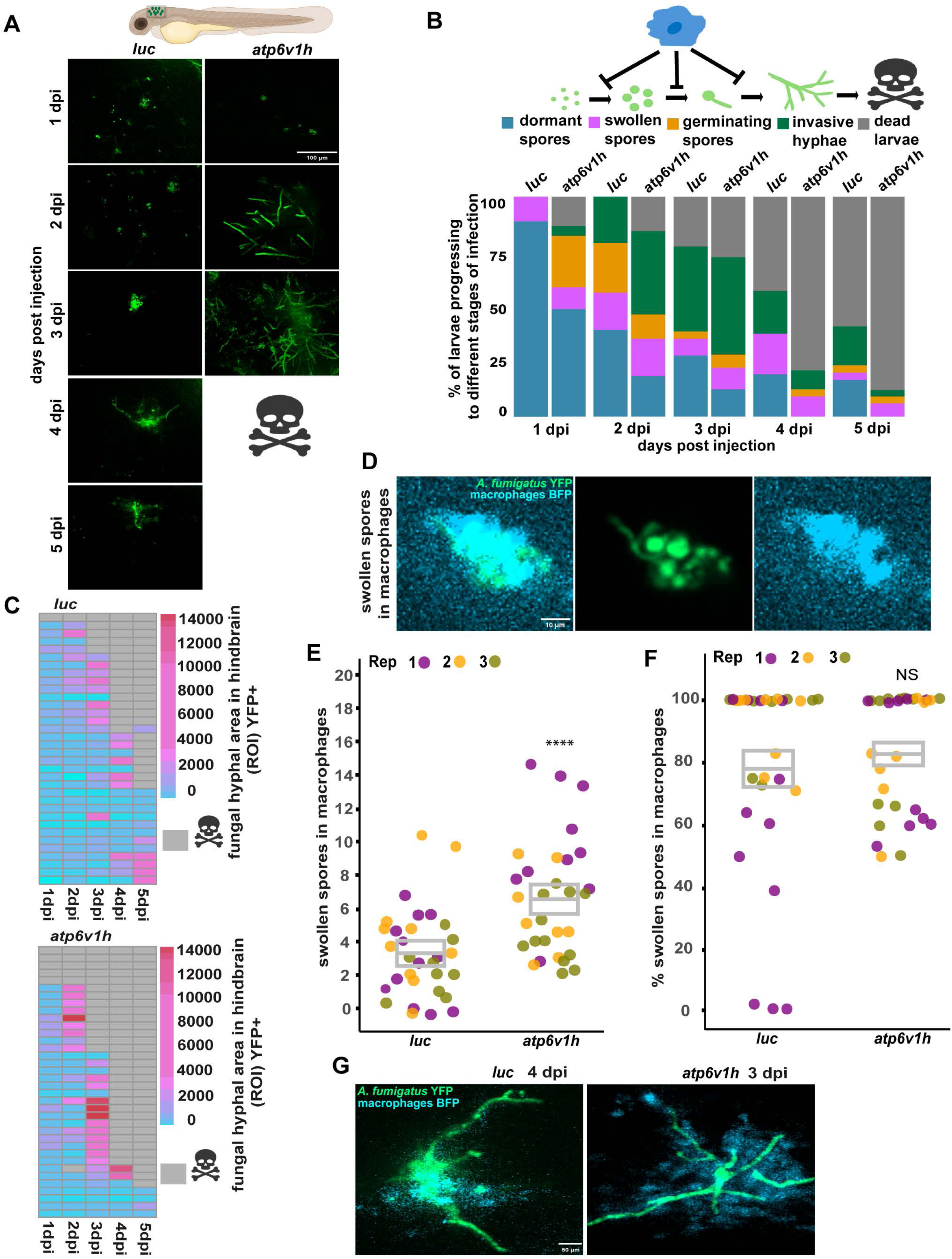
v-ATPase function promotes macrophage control of spore swelling and fungal development. *atp6v1h* crispants and *luc* controls in a neutrophil-defective background (*mpx:rac2D57N*) with fluorescently-labeled macrophages (*mfap4:BFP*) were infected with *Af*293-derived YFP-expressing *A. fumigatus* spores (TBK1.1) and imaged from 1 to 5 dpi. (A) Representative confocal images. Scale bar: 100 µm. (B) Cumulative percentage of larvae showing progression of fungal development into five stages: dormant spores, swollen spores, germinating spores, invasive hyphal growth, and death of larvae. (C) YFP-positive fungal area in the hindbrain was measured per larvae and displayed as a heat map with gray cells indicating larval death. Each row represents a single larva across 5 days of infection. (D) Example images of swollen spores in macrophages at 3 dpi. Scale bar: 10 µm. (E-F) Number of swollen spores inside macrophages per larva (E) and percentage of swollen spores that are inside of macrophages (F) was quantified. Each dot represents an individual larva, with colors indicating separate experimental replicates. *P* values were calculated by emmeans and ANOVA, gray boxes represent the emmeans ± SEM. (G) Example images of recruitment of macrophages to hyphae. Scale bar: 50 µm.

As significantly fewer spores are phagocytosed by macrophages in *atp6v1h* crispant larvae (Fig 4C), one explanation for this increased hyphal growth is that non-phagocytosed spores swell and germinate. However, spores can also swell and germinate from within macrophages (Fig 6D). To assess the role of *atp6v1h* in macrophage-mediated control of spore germination, we quantified the number of swollen spores within macrophages at 2 dpi, finding that *atp6v1h* crispants have a significantly higher number of swollen spores in macrophages compared to *luc* controls (Fig 6E). Additionally, the overall percentages of swollen spores that are found within macrophages as opposed to extracellularly was not significantly different in *atp6v1h* crispants compared to *luc* controls (Fig 6F). These data suggest that the increased fungal development observed in *atp6v1h* crispant larvae is not solely due to germination and growth of spores that are never taken up by macrophages, but instead that Atp6v1h-deficient macrophages are not able to effectively inhibit spore swelling and germination.

### Macrophage-mediated control of post-germination hyphal growth is decreased in v-ATPase deficient larvae

During live daily imaging of fungal development, we noticed that at later stages of infection, high numbers of macrophages accumulated at the infection site even in *atpv1h* crispants (Fig 6G). As we have previously shown that macrophages can play a role in controlling fungal growth post-germination (Tanner & Rosowski, 2024), we wondered whether v-ATPase deficiency affects the ability of macrophages to respond to and control fungal growth post-germination. To specifically investigate this question, we infected neutrophil-defective *atp6v1h* crispant and *luc* control larvae that had fluorescently-labeled macrophages (*mfap4:BFP*) with YFP-expressing *Af*293-derived spores and then screened larvae at 2 or 3 dpi to select larvae that already had developing hyphae. We then performed timelapse imaging for 12 hours to track macrophage numbers and hyphal growth post-germination (Video 2A, B, Fig 7A). Normalizing fluorescent-positive hyphal area to the start of imaging, *atp6v1h* crispants exhibited significantly increased hyphal growth compared to *luc* controls at 12 hours post imaging (Fig 7B). However, this uncontrolled growth is not due to a lack of macrophage recruitment to hyphae, as the normalized macrophage area at the infection site is also increased significantly in *atp6v1h* crispant larvae compared to *luc* controls (Fig 7C). In fact, plotting the relationship between fungal area and macrophage area across the timelapse, macrophage numbers are relatively stable in *luc* control larvae while macrophage numbers are positively correlated with fungal area in *atp6v1h* crispants (Fig 7D). These data demonstrate that macrophages in *atp6v1h* crispants respond to fungal hyphal growth but fail to contain it, whereas *luc* controls exhibit more effective fungal control, resulting in overall lower hyphal growth. These data suggest that v-ATPase function is required for macrophages to control hyphal growth post-germination, independent of any defect in macrophage numbers or recruitment.

**Figure 7:**
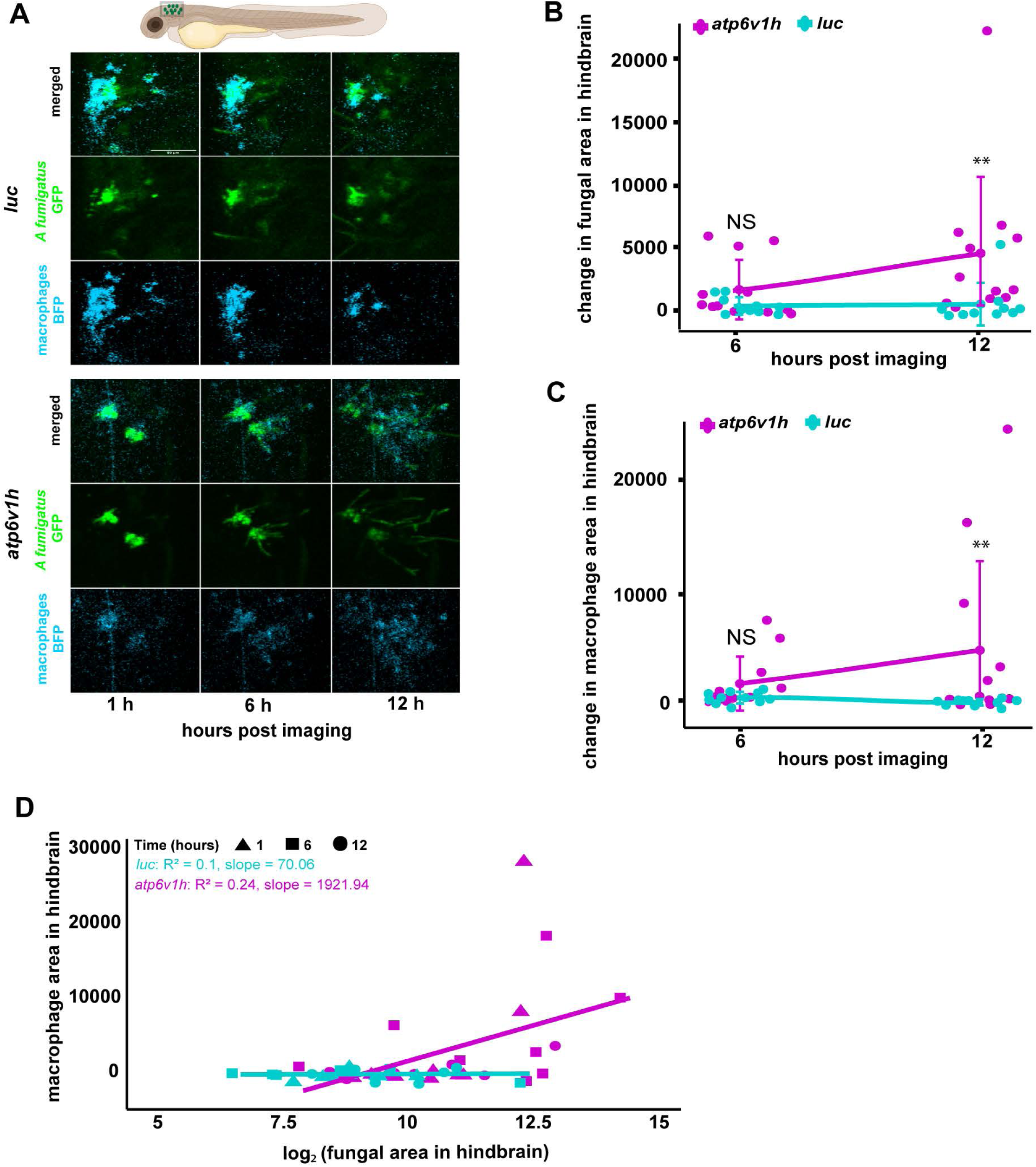
Macrophage-mediated control of hyphal growth following germination is impaired in v-ATPase-deficient larvae. *atp6v1h* crispants and *luc* controls in a neutrophil-defective background (*mpx:rac2D57N*) with fluorescently-labeled macrophages (*mfap4:BFP*) were infected with *Af*293-derived YFP-expressing *A. fumigatus* spores (TBK1.1). At 2 and 3 dpi, larvae were screened for the presence of germination and timelapse imaged for 12 hours. (A) Example images. Scale bar: 50 µm. (B, C) YFP+ hyphal area (B) and BFP+ macrophage area (C) was quantified and normalized to values at 1 hour post-imaging start. (D) Scatter plot of log2 hyphal area versus macrophage area for individual larvae across the timelapse, with slopes of best fit lines indicated. Each dot represents an individual larva, data is pooled from three independent replicates. *P* values were calculated by emmeans and ANOVA, bars represent the emmeans ± SEM.

### v-ATPase deficiency phenocopies increased fungal germination and invasive growth in macrophage-deficient larvae

Our results demonstrate that v-ATPase function is required for multiple aspects of macrophage biology, including total macrophage numbers, acidification of phagocytosed spores, inhibition of spore swelling, and control of hyphal growth post-germination. In *atp6v1h* crispant larvae, *A. fumigatus* spores germinate as early as 1 dpi, which is a similar timeline to what we have previously observed in *irf8* mutant larvae that completely lack macrophages ^8^. We therefore wondered if macrophages that lack functional v-ATPase have any significant protective effect compared to a complete lack of macrophages. To directly compare these two scenarios, we in-crossed *irf8*^+/-^ adults to generate *irf8*^+/+^, *irf8*^+/-^, and *irf8*^-/-^ embryos. We generated *atp6v1h* or *luc* control crispants in these progeny and infected larvae with *Af*293-derived YFP-expressing spores (Fig 8A). We performed live imaging at 1 and 2 dpi and quantified fungal germination, progression to invasive growth, and total fluorescent-positive fungal area. Both *luc irf8*^+/+^ and *luc irf8*^+/-^ larvae exhibited minimal germination and progression of germination to invasive hyphal growth (Fig 8B, C). As previously observed, in control *irf8*^-/-^ larvae that lack macrophages, >50% of larvae experience germination by 1 dpi (Fig 8B). In *atp6v1h* crispants in a wild-type *irf8*^+/+^ background, only ∼20% of larvae experience germination at 1 dpi, but by 2 dpi, this percentage increases to >50%, matching the percentage of *irf8^-/-^*larvae with germination. The percentage of larvae that have germination that progresses to invasive hyphae is also similar in *luc irf8^-/-^*and *atp6v1h irf8^+/+^* larvae (Fig 8C). Similar results were also seen from quantification of total fluorescent-positive fungal area, with comparable fungal area in *luc irf8^-/-^* and *atp6v1h irf8^+/+^* larvae (Fig 8D). These data suggest that the phenotype of larvae that lack v-ATPase function is similar to that of larvae that completely lack macrophages.

**Figure 8:**
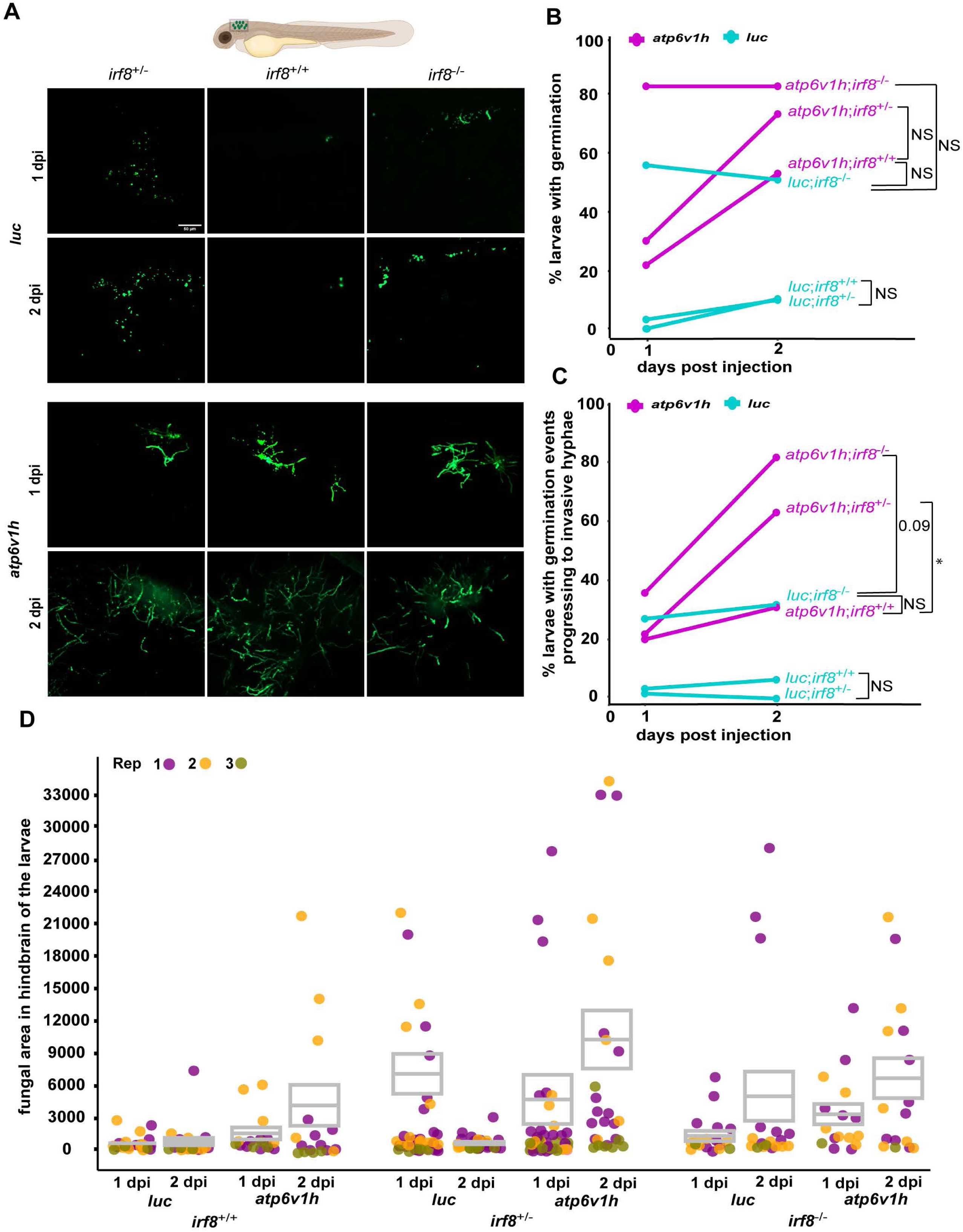
v-ATPase- and macrophage-deficient larvae have similar control of fungal germination and growth. *atp6v1h* crispants and *luc* controls in *irf8*^+/+^, *irf8*^+/-^ and *irf8*^-/-^ backgrounds were infected with Af293-derived YFP-expressing *A. fumigatus* spores (TBK1.1) and imaged at 1 and 2 dpi. (A) Representative images. Scale bar: 50 µm. (B, C) Percentage of larvae with spore germination (B) and percentage of larvae with germination events that progressed to invasive hyphae (C) was quantified. *P* values were calculated by Fisher’s exact test. (D) YFP+ fungal area was quantified. Each dot represents an individual larva, with colors indicating separate experimental replicates. *P* values were calculated by emmeans and ANOVA, and bars represent the emmeans ± SEM.

When we generated *atp6v1h* crispants in an *irf8^-/-^*or *irf8^+/-^* background, an even higher percentage of these larvae experienced germination and germination progressing to invasive hyphae than either *atp6v1h* crispants alone or *irf8* mutants alone (Fig 8B, C). In fact, *atp6v1h irf8^+/-^* larvae had the highest fungal burden of all conditions at 2 dpi (Fig 8D). Heterozygotes in *irf8* still develop macrophages ^59^ and a similar percentage of *luc irf8^+/-^* larvae experience germination and the progression to invasive hyphae as *luc irf8^+/+^*larvae (Fig 8B, C). However, the increase in appearance of germination and progression to invasive hyphal growth observed when the *irf8^+/-^* background is combined with a deficiency in v-ATPase suggests that *irf8* heterozygosity leads to a partial loss of function that compounds the *atp6v1h* crispant defect in macrophage function. In macrophage-deficient *luc irf8^-/-^*larvae, the progression of germination to invasive hyphae is controlled by neutrophils, and neutrophil activity explains why some germination is cleared in these larvae from 1 to 2 dpi (Fig 8B) and why only 25-30% of these larvae progress to invasive hyphae (Fig 8C). A higher percentage (80%) of *atp6v1h irf8^-/-^* larvae experience germination at 1 dpi and this germination is maintained at 2 dpi (Fig 8B). Additionally, ∼80% of *atp6v1h irf8^-/-^* larvae progress to invasive hyphae at 2 dpi (Fig 8C). These data suggest that *atp6v1h* also impairs neutrophil function against hyphae, leading to decreased fungal clearance in the absence of macrophages. Overall, data from this experiment demonstrate that while v-ATPase deficiency phenocopies the increased fungal germination and invasive growth observed in macrophage-deficient larvae, v-ATPase deficient macrophages still have some ability to control fungal germination and growth that can be compromised by a partial loss of function in Irf8 observed in *irf8* heterozygotes. Additionally, v-ATPase likely has additional roles in the ability of neutrophils to control and kill *A. fumigatus* hyphae.

## Discussion

In immunocompetent individuals, alveolar macrophages serve as a primary defense against *A. fumigatus* by phagocytosing spores ^60^. However, the precise trafficking and killing mechanisms that macrophages use to control spores in vivo remain unclear. Here, we used a larval zebrafish host model to examine the role of the v-ATPase complex in macrophage-mediated control of *A. fumigatus*. Surprisingly, we found that while the disruption of v-ATPase activity compromises many aspects of macrophage-mediated fungal defense, v-ATPase-mediated phagosomal acidification is not necessary for spore killing. Instead, v-ATPase-deficient larvae succumb to infection due to loss of control of germination and hyphal growth, demonstrating that spore killing and inhibition of germination can be separable macrophage functions.

A complication in our study is that *atp6v1h* crispant larvae have significantly lower overall numbers of macrophages. One reason for lowered cell numbers is that more macrophages undergo apoptosis. v-ATPase deficiency can compromise lysosomal integrity, leading to lysosomal membrane permeabilization and release of catabolic enzymes into the cytosol which can cleave pro-apoptotic proteins like Bid, initiating mitochondrial cytochrome c release and caspase activation ^61–63^. Atp6v1h deficiency likely resembles lysosomal storage disorders, a group of inherited metabolic diseases characterized by accumulation of undigested macromolecules within lysosomes due to deficiency of transport proteins which particularly affects macrophages ^64–66^. We do report a significantly higher number of TUNEL+ apoptotic macrophage nuclei in the caudal hematopoietic tissue of *atp6v1h* crispant larvae. However, these numbers increase from ∼1 to ∼3.5 per CHT, which cannot fully explain the change from ∼225 to ∼150 total macrophages that we find across whole larvae. It is possible that macrophage apoptosis increases further in these larvae under the stress of infection ^17^.

Impaired myeloid development likely also contributes to the decreased macrophage and neutrophil numbers in *atp6v1h* crispant larvae. We do find significantly increased apoptosis in surrounding cells compared to macrophages within CHT of *atp6v1h* crispant larvae compared to control larvae. One possible mechanism is disruption of the hematopoietic microenvironment, potentially impairing the survival or function of non-macrophage niche cells essential for myeloid development. Previous studies in zebrafish show v-ATPase activity is essential for the development, proliferation, and function of diverse cell types, including those involved in ciliogenesis, retinal morphogenesis, and sensory organ formation ^67–69^. For example, Knockouts of v-ATPase subunits cause diverse defects: *atp6v1h* knockout impairs bone remodeling and leads to osteopetrosis ^70^, *atp6v1a* causes suppression of acid-secretion from skin, growth retardation, trunk deformation, and loss of internal Ca^2+^ and Na^+^ ^71^, v-ATPase accessory protein Atp6ap1b depletion caused defects in neuromasts and olfactory placodes ^72^, and the mutations in core components like *atp6v1h*, *atp6v1f*, *atp6v1e1*, *atp6v0c*, and *atp6v0d1* elevated levels of apoptotic neurons within their retinas and brains ^67^. Thus, while Atp6v1h deficiency does directly lead to some macrophage apoptosis, impaired lysosomal function and increased cell death in the hematopoietic niche may reflect a broader disruption of tissue homeostasis and myeloid cell development.

Through live imaging and detailed analysis we were able to determine the effect of *atp6v1h* mutation on the function of macrophages that do migrate to the infection site and interact with spores, despite reduced overall macrophage numbers. As expected, the percentage of spores that are inside of macrophages that are in Lysotracker+ acidified compartments is significantly lower in *atp6v1h* crispants, consistent with the requirement of v-ATPase activity to lower intraluminal pH ^18^. Surprisingly, however, we report no difference in the ability of v-ATPase-deficient macrophages to kill *A. fumigatus* spores, either by live-dead staining and live imaging or by CFU counts over time. This finding is consistent with a previous study from our lab finding low correlation between spore acidification and spore killing ^73^. This finding demonstrates that these macrophages can use pH-independent mechanisms such as reactive oxygen species or antimicrobial peptides to compensate for reduced acidification ^74^.

In wild-type larvae, only ∼40% of phagocytosed spores are acidified at 1 dpi. We were also able to visualize v-ATPase localization on macrophage phagosomes inside of a live intact host for the first time, through transgenic expression of fluorescently-tagged Atp6v1h, and we report that only ∼35-50% of spores in macrophages are co-localized with Atp6v1h, matching the percentage of spores that are acidified. These results, and the dependence of macrophages on other killing mechanisms, likely reflect the ability of *A. fumigatus* to actively inhibit phagosomal acidification. *A. fumigatus* urease decreases phagosomal acidification, enhancing spore survival within macrophages ^75^. Spores can also prevent phagosomal trafficking to the lysosome ^15^. For example, DHN-melanin, produced by the polyketide synthase PksP, masks the cell wall, actively inhibiting spore recognition and phagosome–lysosome fusion ^53,76^. In epithelial cells, the heat shock protein HscA of spores can redirect spores into a non-degradative recycling pathway, but it is unclear if this occurs inside of macrophages ^14^. The impaired acidification of *A. fumigatus* spores in macrophages is therefore likely due to a combination of impaired recognition and trafficking to phago-lysosomes and active detoxification of acidic compartments.

Even though rates of spore killing are not affected by v-ATPase deficiency, *atp6v1h* crispants exhibit accelerated spore germination and hyphal growth that leads to host death. This phenotype is not primarily due to increased germination of non-phagocytosed spores, as the percentage of swollen spores that are inside macrophages versus outside macrophages is not significantly different in *atp6v1h* crispant larvae than controls. Instead, more spores inside of macrophages can swell when the v-ATPase complex is defective, indicating that while it is not required for spore killing, phagosomal acidification is required to inhibit spore germination, and the control of spore germination and spore killing are separable functions of macrophages. How acidification controls germination without killing spores is unclear and should be an area of future study. One potential mechanism is the activation of acid hydrolases such as cathepsins B, L, and K, and β-hexosaminidase which depend on an acidic environment for activity and have been shown to directly mediate microbial control ^77–80^.

We also report that v-ATPase activity promotes macrophage control of *A. fumigatus* hyphal growth post-germination. While neutrophils are the primary innate immune cell to control large extracellular pathogen growth such as *Aspergillus* invasive hyphae, we previously found that in neutrophil-defective larvae macrophages do have some ability to inhibit hyphal growth ^73^. While in *atp6v1h* crispant larvae we found decreased macrophage recruitment to spores; in larvae with hyphae we observe higher macrophage recruitment, demonstrating that defects in hyphal control are due to specific defects in macrophage function. It is still unclear what mechanisms macrophages use to control extracellular growth of this pathogen, but we now know that these mechanisms are dependent on both v-ATPase function and the Rac2 GTPase ^73^. Macrophages can generate extracellular traps to control fungi ^81–83^, and it is possible that acid-dependent lysosomal hydrolases are part of these extracellular traps. It is also possible that the v-ATPase does not play a direct function in extracellular fungal control, but instead that macrophages without v-ATPase function have mitochondrial damage, impaired autophagy, or inflammasome activation that make these cells broadly defective ^17^.

We also identify a potential impact of v-ATPase deficiency on neutrophil function against *A. fumigatus* hyphae as disruption of *atp6v1h* exacerbates infection in *irf8* mutant larvae which lack macrophages ^59,84^. While we do observe a modest reduction in total neutrophil counts in wild-type *atp6v1h* crispants, *irf8* mutants generally have an increased number of neutrophils as cells that would normally become macrophages are re-routed to a neutrophil fate, and thus it is unlikely that the increased fungal growth we observe in *irf8*^-/-^ *atp6v1h* crispants is due primarily to an effect of Atp6v1h deficiency on neutrophil numbers. Whether the mechanisms underlying defective macrophage extracellular fungal killing similarly affect neutrophil-mediated responses in *atp6v1h* crispant larvae remains unclear. It has been shown that the v-ATPase is found on neutrophil granules and treatment with Bafilomycin A1 inhibits myeloperoxidase and elastase secretion ^85^, suggesting that altered neutrophil degranulation is one potential effect of Atp6v1h deficiency in neutrophils that could cause decreased neutrophil-mediated fungal control. Whether other neutrophil mechanisms are also affected and if v-ATPase deficiency in other cells like epithelial cells also promotes susceptibility to *A. fumigatus* also remains to be determined.

In conclusion, our study establishes that v-ATPase activity is indispensable for macrophage-driven immunity against *A. fumigatus* in a whole animal. We identify a decoupling of spore killing and inhibition of spore germination as v-ATPase function is only required for the latter. Additionally, we identify a role for the v-ATPase complex in macrophage-mediated extracellular killing as well as a potential role for this complex in neutrophil extracellular killing functions. Moving forward, it will be important to elucidate how v-ATPase function coordinates phagosomal trafficking, activation of acid hydrolases, and extracellular killing for effective pathogen clearance by different cell types in vivo. A deeper understanding of the multifaceted role of the v-ATPase in immunity could pave the way for identification of v-ATPase modulators that promote control of *A. fumigatus* infection and hyphal growth.

## Author contributions

A.R.M.- Conceptualization, Methodology, Formal analysis, Investigation, Writing - Original Draft, Visualization

E.E.R- Conceptualization, Resources, Writing - Review & Editing, Supervision, Funding acquisition

## Acknowledgments

We thank members of the Rosowski Lab for helpful discussions and for assistance with zebrafish care. We thank the Clemson University Aquatic Animal Research Laboratory for providing the microinjection setup used for single-cell embryo injections. We thank Erin Glass, Bailey Naples, and Savini Thrikawala for technical assistance in generation of transgenic zebrafish.

## Funding Acknowledgements

This research was supported by the National Insitutes of Health: the National Institute of Allergy and Infectious Diseases under award number R21AI164363 to E.E.R. and the National Institute of General Medical Sciences under award number R35GM147464 to E.E.R.

## Abbreviations

CHT: Caudal hematopoietic tissue
dpf: days postfertilization
dpi: days postinjection
emmean: estimated marginal mean
GMM: glucose minimal medium
gRNA: guide RNA
HR: hazard ratio
PTU: N-phenylthiourea
TIDE: Tracking of Indels by Decomposition
TUNEL: Terminal deoxynucleotidyl transferase dUTP Nick-End Labeling

## Figure legends

**Suppl Fig 1: CRISPR/Cas9-mediated targeting of *atp6v1h* in zebrafish**

(A) Gene structure of *atp6v1h* from exons 3 to 5 with target sites for two gRNAs and primers used to confirm targeting. (B) Schematic representation of the experimental setup for cloning-free in vitro gRNA synthesis showing the generation of a DNA template containing a targeting sequence fused to the constant region under a T7 promoter. (C) PCR amplification of DNA from 1 dpf embryos injected with gRNAs and Cas9. Each lane corresponds to a PCR reaction with genomic DNA from a single embryo. For F1/R1 and F2/R2 amplification, smearing indicates the mosaic introduction of indels. For F1/R2 amplification, a band is generated only if intervening DNA between gRNA target sites is lost. (D) Histogram plots from TIDE analysis showing CRISPR-Cas9 editing efficiency for *atp6v1h* gRNAs 1 and 2. Red bars indicate statistically significant mutations (*p* < 0.001, R² values shown), while black bars represent non-significant changes. Overall editing efficiencies range from 52.2% to 69.3%, as indicated above each plot.

**Suppl Fig 2: Bafilomycin does not reduce survival of control larvae**

Larvae in a neutrophil-defective background (*mpx:rac2D57N*) were injected with PBS, treated with Bafilomycin A1 (BafA1), or DMSO vehicle control and monitored for survival for 7 days.

**Suppl Video 1A, B: Timelapse imaging of macrophage response after injection**

*atp6v1h* crispants or *luc* control larvae with fluorescently-labeled macrophages (*mfap4:BFP*; *mpeg1:H2B-GFP*) were infected with RFP-expressing *Af*293-derived *A. fumigatus* spores (TBK5.1) and followed by confocal live timelapse imaging from 1-18 hours post injection. Videos comprise max intensity z projections.

**Suppl Fig 3: v-ATPase deficiency impairs overall neutrophil numbers**

*atp6v1h* crispants and *luc* controls with fluorescently-labeled neutrophils (*lyz:H2B-mCherry*) were imaged at 2 days post-fertilization (dpf). Total neutrophil numbers per larva were quantified. Each dot represents an individual larva, with colors indicating separate experimental replicates. *P* values were calculated by emmeans and ANOVA, gray boxes represent the emmeans ± SEM.

**Suppl Fig 4: *atp6v1h* crispants have decreased lysosomal acidification**

(A-B) *atp6v1h* crispants and *luc* controls with fluorescently-labeled macrophages (*mfap4:BFP*) were injected with *Af*293-derived YFP-expressing *A. fumigatus* spores (TBK1.1) with AlexaFluor633 conjugated to the cell wall. At 1 dpi, larvae were stained with Lysotracker Red dye for 1 hour and then confocal imaged. Total lysotracker+ area in macrophages (A) and lysotracker+ area normalized to the total macrophage area (B) was quantified. Each dot represents an individual larva, with colors indicating separate experimental replicates. *P* values were calculated by emmeans and ANOVA, and bars represent the emmeans ± SEM.

**Suppl Video 2A, B: Timelapse imaging of macrophage responses to hyphae**

*atp6v1h* crispants and *luc* controls in a neutrophil-defective background (*mpx:rac2D57N*) with fluorescently-labeled macrophages (*mfap4:BFP*) were infected with *Af*293-derived YFP-expressing *A. fumigatus* spores (TBK1.1). At 2 and 3 dpi, larvae were screened for the presence of germination and timelapse imaged for 12 hours. Scale bar: 100 µm.

