## Supplementary figures and images for "Macrophage vacuolar ATPase (v-ATPase) function controls *Aspergillus fumigatus* germination and hyphal growth independent of spore killing"

### Supplemental Figures 1-4

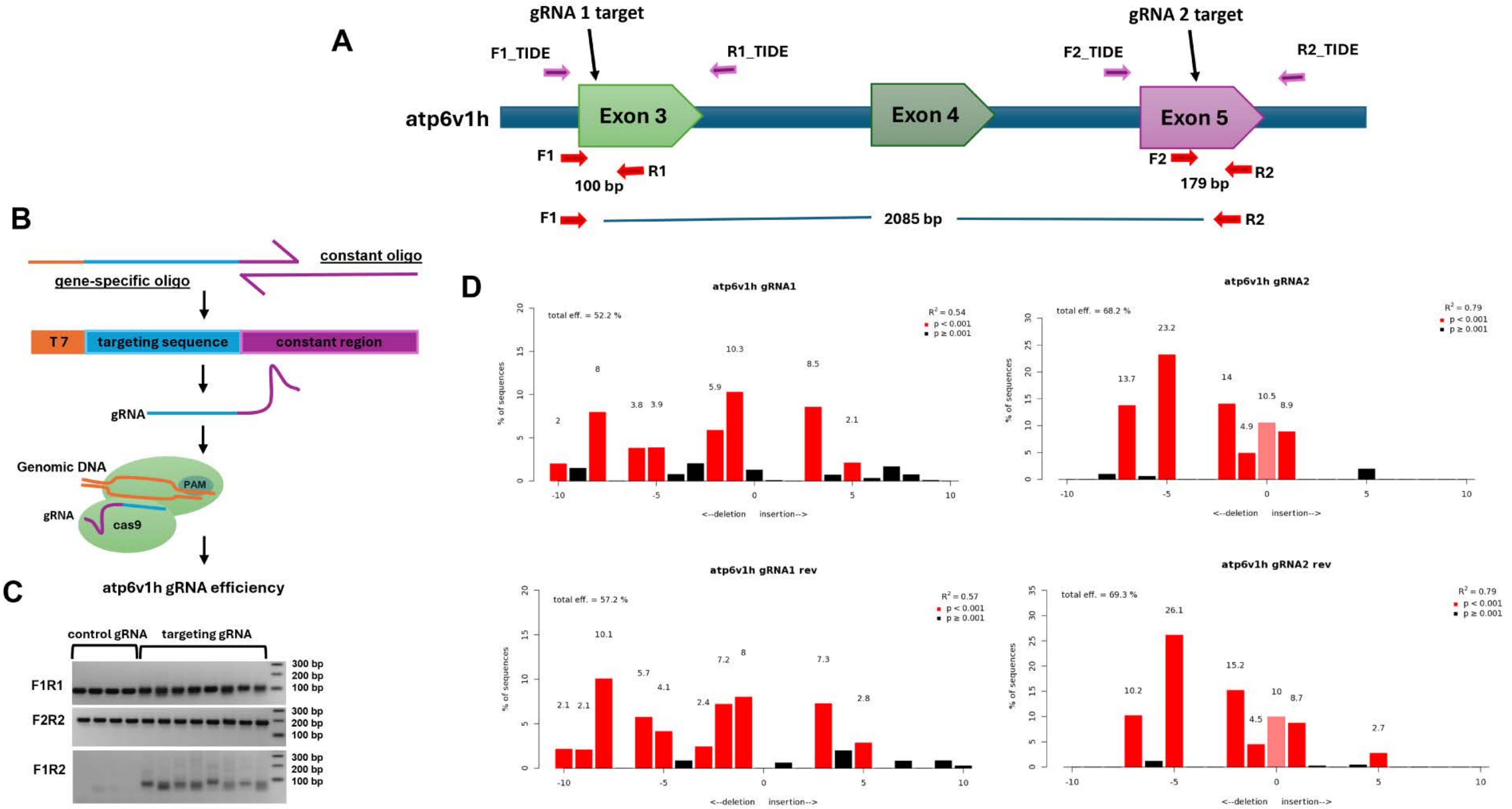

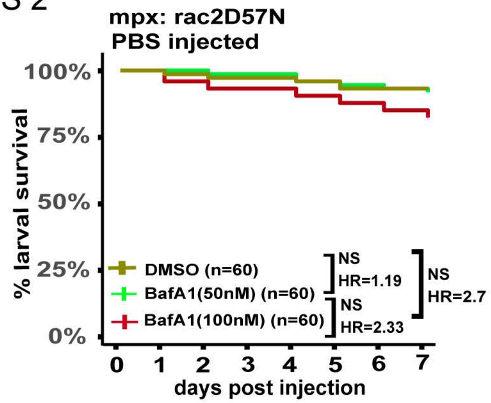

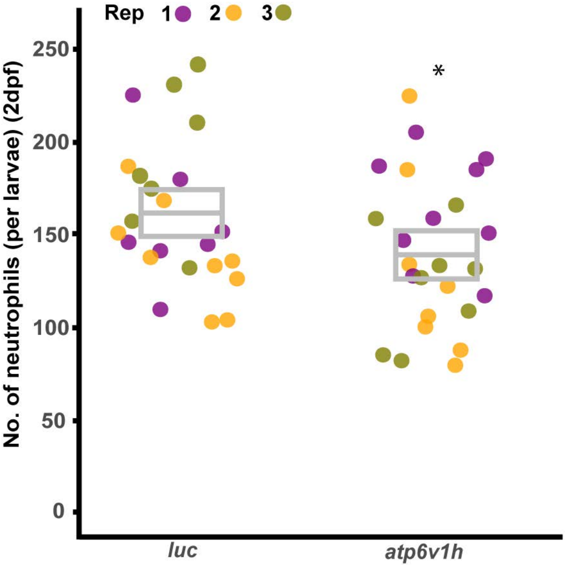

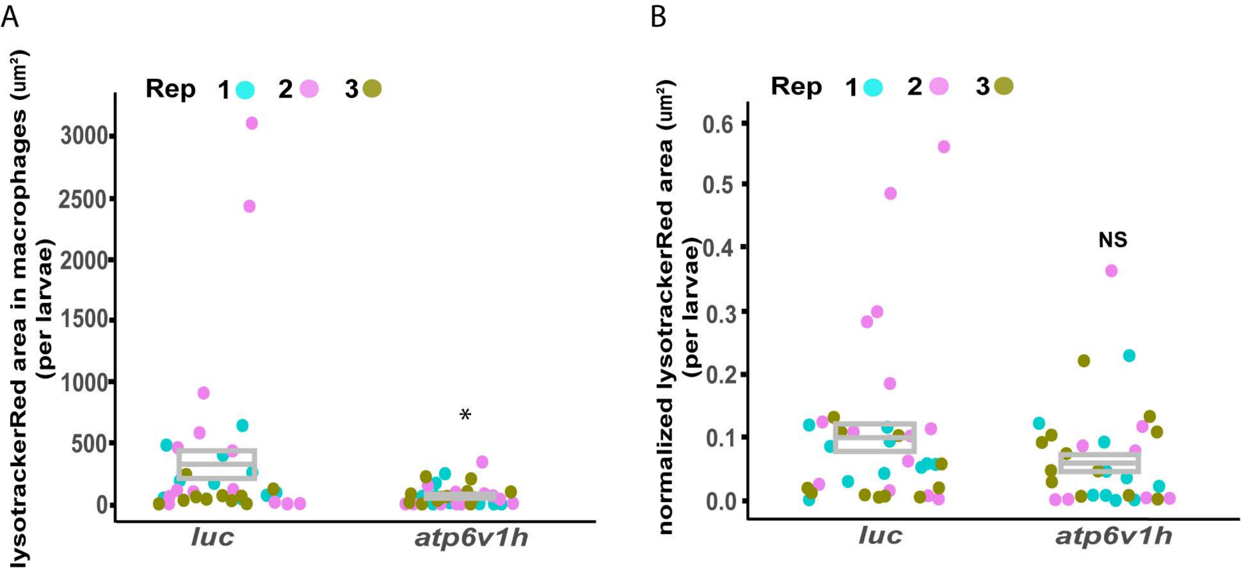
